# Titin as a mechanical damper: Balancing Stability and longevity through inter-domain linker design

**DOI:** 10.1101/2025.03.03.641210

**Authors:** Pritam Saha, Tanuja Joshi, Gaurav K. Bhati, Akriti Adarsh, Deepali Bisht, Sabyasachi Rakshit

## Abstract

Titin, a giant protein (∼3-4 MDa), functions as a molecular-spring to regulate muscle elasticity. More than 90% of Titin is composed of domains that absorb mechanical energy and undergo stochastic unfolding-refolding under tension (∼tens of pN). These domains are connected in tandem by interdomain linkers (IDLs), which constitute less than 10% of the total mass. Despite their small genomic footprint, bioinformatics mapping suggests that IDLs have an outsized impact on protein mechanics, potentially contributing to disease pathology. Using magnetic tweezers, here we examine how linkers influence mechano-response of domains to constant and oscillatory forces. We found that short linkers limit interdomain movement and promote first-order cooperative folding transitions of domains. In contrast, long flexible linkers induce creep-like deformations interspersed with sharp, stepwise transitions. Surprisingly, linkers that improve domain-stability resist unfolding under constant pulling forces, but lose power retention faster under oscillatory forces. Our findings reveal a trade-off between mechanical stability and energy retention in titin, a key muscle protein. These insights offer new design principles for mechano-responsive protein engineering.

**Teaser:** Tiny linkers fine-tune how bulky domains in titin respond to force.

## Introduction

In living organisms, the mechanobiome represents a complex system that processes various mechanical inputs from the surrounding environment(1). These mechanical inputs can differ significantly in magnitude, frequency, and duration, encompassing everything from subtle cues to substantial forces(2). Subtle mechanical cues are crucial for essential cellular and physiological functions(3), influencing processes such as cell growth(4), migration(5), tissue homeostasis(6), and mechanotransduction(7). Conversely, more intense mechanical forces can act as detrimental factors that threaten cellular integrity and function(8), leading to conditions like atherosclerosis or mechanical fatigue. Experimental studies have revealed how cells sense and respond to mechanical stimuli(9), highlighting the essential role of mechanobiome in health and adaptation(10).

Specialized multidomain design of mechanical proteins and protein complexes are fundamental to the mechanobiome, balancing beneficial mechanical cues that sustain cellular functions against harmful forces that threaten cellular integrity(11). These proteins typically exhibit a characteristic long and modular architecture, comprising multiple domains or repeats connected at their N- and C-termini(12). This structure enables multidomain proteins (MDPs) to selectively filter mechanical signals, distinguishing functional cues from damaging assaults(13), thereby preserving the stability and functionality of the mechanobiome.

Domains, the smallest autonomous folding units in proteins, play a crucial role in MDPs(14). Occupying a substantial portion of their volume, domains are considered as primary responders to mechanical stimuli in physiology(14). Subsequently, as barrage of studies, experimentally(15) and computationally(16), have been performed by exposing domains under mechanical tension, revealing their roles as energy storage units(17), power generators(17), shock absorbers(18), and viscoelastic springs(19). Such studies often focus on their unfolding and refolding dynamics in response to mechanical inputs(13, 20), highlighting their functional capabilities.

However, a critical question remains: how do individual domains within an MDP fine-tune their mechanical properties to adapt to the complex force environments of the mechanobiome, despite their structural similarity with neighboring domains in MDP? For example, Titin, the largest human mechanical protein, consists of 244 domains primarily of the Ig-like or FnIII-like superfamilies, each with a similar β-strand rich architecture(21, 22)(Figure 1a). Understanding how these repetitive structures achieve functional diversity to support the dynamic demands of the mechanobiome presents a significant challenge in the field.

**Fig 1.**
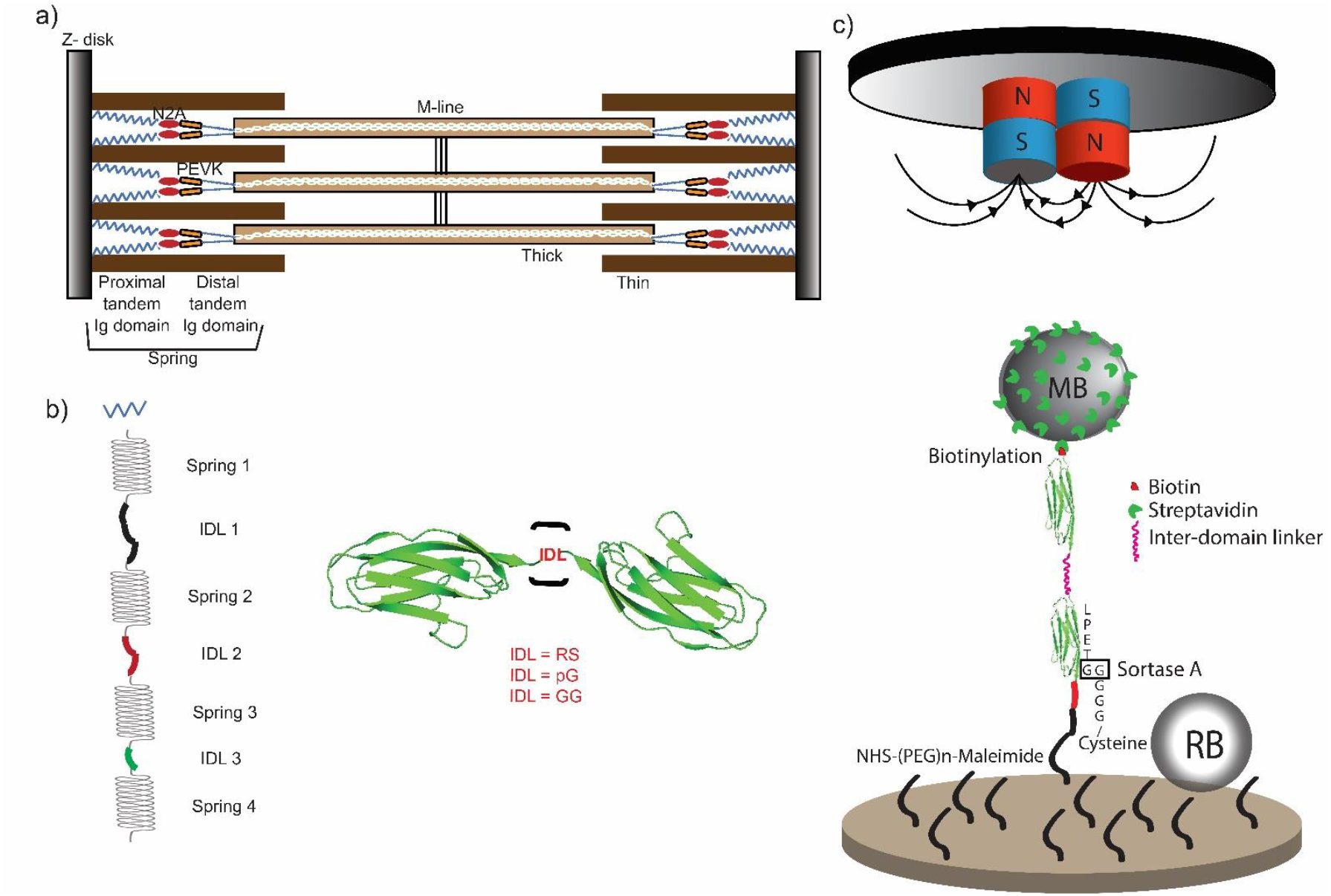
Titin and interdomain linker models and their tethering in MT. a) A schematic depiction of titin protein chain along with the thin and thick filaments in striated muscle sarcomere. Only two titin molecules per half thick filament are shown for simplicity. b) A cartoon representation of three models of two titin domain joined with three different IDLs (RS, GG and GGGSGGGG (pG)). c) A schematic representation of attachment setup of I27 dimer variants in magnetic tweezers. The interdomain linker in red color is the varying parameter here. The I27 dimer (in Green) is covalently attached to the coverslip and robustly attached to streptavidin modified M-270 paramagnetic Dynabeads (MB) (in light brown) using biotinylation. Reference beads (RB) (in Grey) are adsorbed to the surface.

In contrast to domains, interdomain linkers (IDLs) display significant variability in length, physico-chemical properties, and structures(23). Far from being mere spacers, IDLs play a pivotal role in shaping the mechanical stability(24) and functionality(25) of MDPs. Recently, it is reported that IDLs influence the thermodynamic stability(24), kinetics, and mechanical resilience of adjacent repeats of 94^th^ immunoglobulin domain (commonly referred as I27)(26). For example, flexible glycine-rich linkers reduce unfolding forces, while stiffer arginine-serine (RS) linkers enhance domain stability under mechanical stress(27). These findings suggest that identical domains, when connected by different IDLs, may exhibit distinct mechanical responses, positioning IDLs as key modulators of mechanics in MDPs.

To address critical questions in the field, we asked: How do variations in interdomain linker properties modulate the balance between energy retention and mechanical stability in multidomain proteins? Can differences in linker rigidity and chemical composition lead to distinct mechano-responses under constant versus oscillatory forces? Our bioinformatics analysis had revealed a significantly higher prevalence of mutations in interdomain linkers (IDLs) compared to domains, despite the substantially smaller number of amino acids in IDLs (Supplementary table 1). To explore these questions, we engineered I27 dimer variants incorporating a short, rigid linker (−GG-), a previously studied -RS-linker, and a flexible, chemically inert linker (GGGSGGGG, or pG) that preserves domain identity (Figure 1b). We then subjected these constructs to a range of forces using both experimental and computational methods. This approach allowed us to explore how the interplay between domains and their linkers influences the mechanical behavior of multidomain proteins. Our findings highlight a critical balance: the trade-off between optimal energy retention for extended durations and the mechanical stability required for essential biological processes.

## Results

In line with the reported studies(24), our ensemble thermodynamic measurements using fluorescence spectroscopy and far-UV circular dichroism too indicated that the chemical and thermal stability of the I27 dimers follow the order: RS > pG > GG (SF 1).

To map the comparative nano-mechanics of the dimer variants in response to both constant and oscillatory force inputs, we then performed single-molecule force spectroscopy measurements utilizing our laboratory-built magnetic tweezers (MT)(20) (SF 2).

### Single-molecule tethering of titin for Magnetic Tweezers (MT) experiments

For MT experiments, the linker variants (Avitag-I27-linker-I27-LPETGG) were covalently attached to the coverslip at its C-terminus via Sortase A-mediated ligation (see Methods), while the N-terminus was linked to a paramagnetic bead (d ∼ 2.8 μm) using biotin-streptavidin interactions (Figure 1c). A detailed description of surface modifications and protein attachment protocol is written in the Method section(20). A resting force of 5.0 pN was applied to eliminate non-specific interactions between the bead and the coverslip. Non-magnetic beads were used to correct for drift during measurements. Extension and lifetime of unfolding and refolding events were analyzed using Autostepfinder(28), a GUI based on the mean standard deviation model.

To optimize single-molecule tethering in MT experiments, we standardized surface preparation by systematically varying APTES concentrations (0.5–5%). Lower APTES concentrations (0.5–1%) effectively minimized multi-tether events by reducing immobilization density. Covalent attachment mediated by Sortase A further improved the efficiency of single-molecule tethering while eliminating non-specific interactions. Optimization in tethering was validated by monitoring the entropic extension (>20 nm) under increasing forces during force clamp experiments(20), followed by the stepwise unfolding of I27 domains at high forces (>100 pN)(20). The relation of the unfolding lengths of I27 domain with clamping forces conformed to the worm-like chain (WLC) model, further confirming that unfolding events arose from specific molecular interactions at the single-molecule level (SF 3). We next performed two types of force-clamp experiments, *constant* force-clamp experiments and *oscillatory* force-clamp experiments, as discussed below respectively.

I27 of Titin protein possesses high mechano-stability. To identify the clamping forces that unfold domains in our MT, we first clamped all linker variants to 85 pN and rarely observed unfolding events for GG and pG variants even after prolonged clamps of 120 s. RS-variant did not show any unfolding feature at 85 pN within the clamping duration of 120 s, indicating RS-linker variant as the most resistant to tensile forces than the other variants. Thus, to facilitate unfolding events in real-time for all variants, we performed our *constant* force-clamp experiments at relatively higher forces of 103.8, 113.6, and 124.2 pN, respectively with clamp times of 120 s, 90 s, and 60 s, respectively. The forces we calculated from the magnet positions of 3.65 mm, 3.55 mm, and 3.46 mm away from the surface, respectively. For refolding, we first clamped all variants to the highest force of 124.2 pN for 15 s to ensure complete unfolding of domains and then, quenched to 5.0 pN, 7.0 pN, and 8.4 pN.

### Folding dynamics of linker variants from *constant* force-clamp measurements

Figure 1c shows representative force-clamp curves for I27-linker-I27, illustrating their unfolding and refolding behavior under tension. In single-molecule force spectroscopy, unfolding-folding transitions typically involve an initial entropic extension followed by step-like jumps in the contour lengths of domains. However, in our experiments, we observed two distinct types of contour length changes during force-clamp measurements across all linker variants: gradual creep-like changes and more pronounced step-like jumps (highlighted in Figure 3a). Notably, while these transitions occur in tandem, their sequence of occurrence is random.

The step-like jumps, corresponding to transitions between folded and unfolded states, are characteristic of first-order cooperative phase transitions. In contrast, the gradual creep-like changes refer to second-order phase transitions, where the creep rate reflects the degree of cooperativity in the transition. The total extension in contour length, accounting for both step-wise and creep-like changes, aligns with the predicted contour length (Lc) of the I27 dimer (Lc ∼ 25 nm)(20, 29). This finding suggests that the I27 domain in the linker variants can adopt two distinct conformations: one that undergoes a rapid, highly cooperative first-order transition and another that experiences a slower, second-order transition. Although creep-like transitions in protein folding are rarely observed in force-spectroscopy, similar phenomena occur in materials where grain-boundary sliding allows slow deformation under a sustained load. We propose that this creep-like behavior arises from transitions among energetically closely situated microstates within the force-induced folding and unfolding pathways during subdomain sliding. However, the existence of these microstates remains unresolved in our measurements.

Below, we analyzed both step-like jumps and creep-like transitions separately.

### Step-wise instantaneous, cooperative unfolding show variation with different IDLs

From the stepwise changes in bead height, we estimated contour length changes (ΔLc) during step-like unfolding and refolding, and dynamics were analyzed through dwell-time analysis (Figure 2a, see Methods). Data were collected from at least 20 distinct beads per linker variant, with multiple unfolding-refolding cycles performed for each bead.

**Fig 2:**
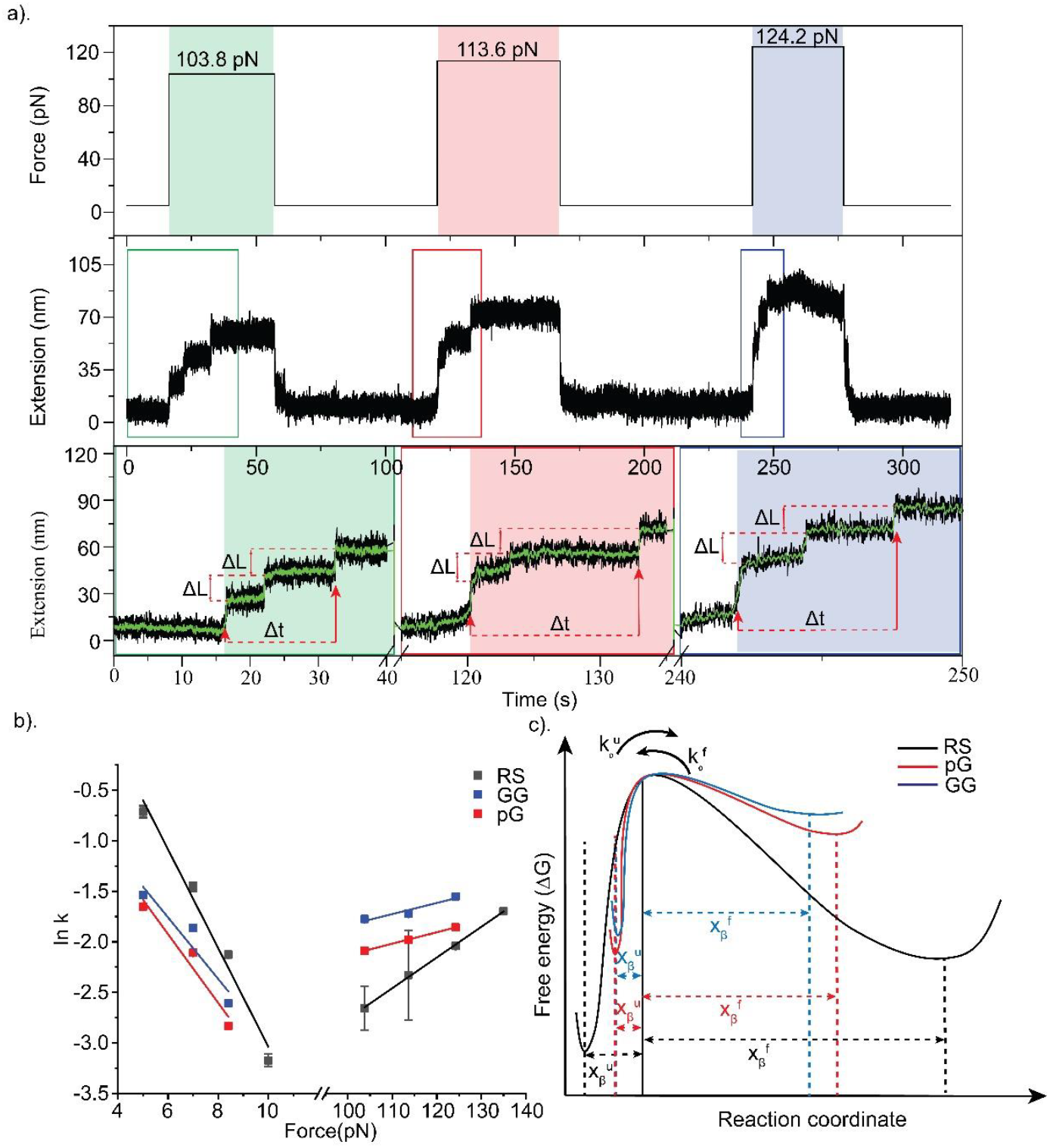
Folding dynamics of three protein models under tensile force. a) A representative extension versus time trace of (I27)2 Linker model at three clamping forces 103.8 pN, 113.6 pN and 124.2 pN. With increase in force, we observed a faster unfolding fingerprint for I27 dimer. After 60s of force clamp, force is quenched to 5 pN for about 45 seconds allowing complete refolding. The highlighted box is zoomed in depiction of the unfolding fingerprint of two domains. The green box is resulting from 103.8 pN, the red box is resulting from 113.6 pN and the blue box is resulting from 124.2 pN. The time-trace part which is under the clamping force has been highlighted with a background shade, respectively. The black data points represent the raw data collected at 200 Hz and the green data points between the black data points are the result of 10 points adjacent to the averaging smoothing filter. Contour lengths and dwell-time are collected from the step-fitting using auto-stepfinder. b) Force-induced unfolding (103.8, 113.6 and 124.2 pN) and refolding (5, 7 and 8.4 pN) rate data for RS (black), pG (red) and GG (blue) linker variants are shown in the Chevron plot. The corresponding solid lines are the fit to the Bell’s equation (ln k_*f*_ = ln k_*0*_ + F * (X*β* / k_*B*_. *T*)). c) Representative 1D potential-energy profiles for all three variants, relatively scaled according to the kinetic parameters (Supplementary Table 2) obtained from the Chevron plot.

ΔLc distributions were consistent across linker variants except for RS at the lowest clamping force (103.8 pN) (SF 3). For RS at this force, a bi-nominal distribution emerged, with peaks at 11.5 ± 0.3 nm and 20.6 ± 0.6 nm. At higher forces, these distributions merged into a single peak (23.1 ± 0.6 nm) (SF 3). While the distinct low-force distribution for RS remains unclear, we propose that the domain-linker interactions modify the force propagation paths in RS variant than the inert linker variants. The domain-linker interactions thus redefine the domain-structures and mediate unique unfolding pathway for RS-variant. At high forces, single-molecule force-clamp spectroscopy of I27 octamers with RS linker consistently showed a single ΔLc distribution with a peak at 23.1 ± 0.6 nm.

Dwell-time distributions for unfolding and refolding were independently fitted to exponential decay functions, providing lifetimes for native and denatured states at respective clamping forces (see Methods, SF 4-5). Similar to ΔLc, the dwell-time distribution of unfolding for I27-RS-I27 exhibited bi-exponential decay at low forces, and converged to a single exponential at higher forces (Figure 2d-f). For the unfolding of GG and pG linkers, the dwell-time distributions follow single exponential decays for all forces. Dwell-time distributions of refolding for all linker variants followed single-exponential decays at all forces. We next fitted the force-lifetime curves to Bell’s model(30), and estimated the intrinsic kinetic parameters such as the zero-force folding 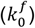 and unfolding rates 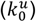 and the distances to the transition states from folded states 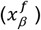 and unfolded states 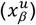 (Supplementary table 2) (SF 4-5). As expected, I27-RS-I27 among the variants, showed the longest native-state lifetime 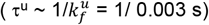, followed by pG (1/0.038 s) and least for GG (1/0.053 s) (Supplementary table 3). We measured the longest 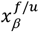 for RS 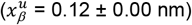 followed by pG 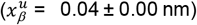 and GG 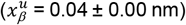. GG displayed the lowest mechanical stability, despite sequence similarity with pG and comparable length to RS. Overall, these results align with reported chemical and thermal stability trends: RS > pG > GG. While the unfolding rates showed significant differences for all three variants, refolding rates were comparable for GG 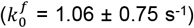 and pG 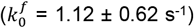 (Figure 2g-i, SI Table 3). RS refolded fastest among the variants 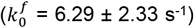. We hypothesize that the interdomain interactions in RS may act as folding nucleators. RS also showed the longest 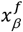 value of -2.00 ± 0.19 nm, while GG had the shortest 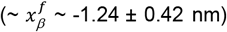, indicating that RS experiences the most pronounced force-induced modifications in the folding free-energy barrier.

### Creep-like gradual, less cooperative unfolding show variation with different IDLs

Creep-like phenomena were observed in a substantial fraction of force-clamp experiments: 37.5%, 34.6%, and 40.4% for RS at clamping forces of 103.8 pN, 113.6 pN, and 124.2 pN, respectively; 30.0%, 30.8%, and 35.3% for GG; and significantly higher rates of 73.0%, 62.2%, and 58.5% for pG at equivalent forces (Figure 3b). Similar trends were evident during the refolding phase post-force quenching, highlighting the reversibility of creep dynamics across conditions (Figure 3e). Creep events during refolding were recorded as 3.8%, 24.5%, and 26.0% for RS at 5 pN, 7 pN, and 8.4 pN; 5.9%, 29.3%, and 26.7% for GG; and notably higher at 6.5%, 47.0%, and 38.5% for pG under equivalent quenching forces (Figure 3f).

**Fig 3:**
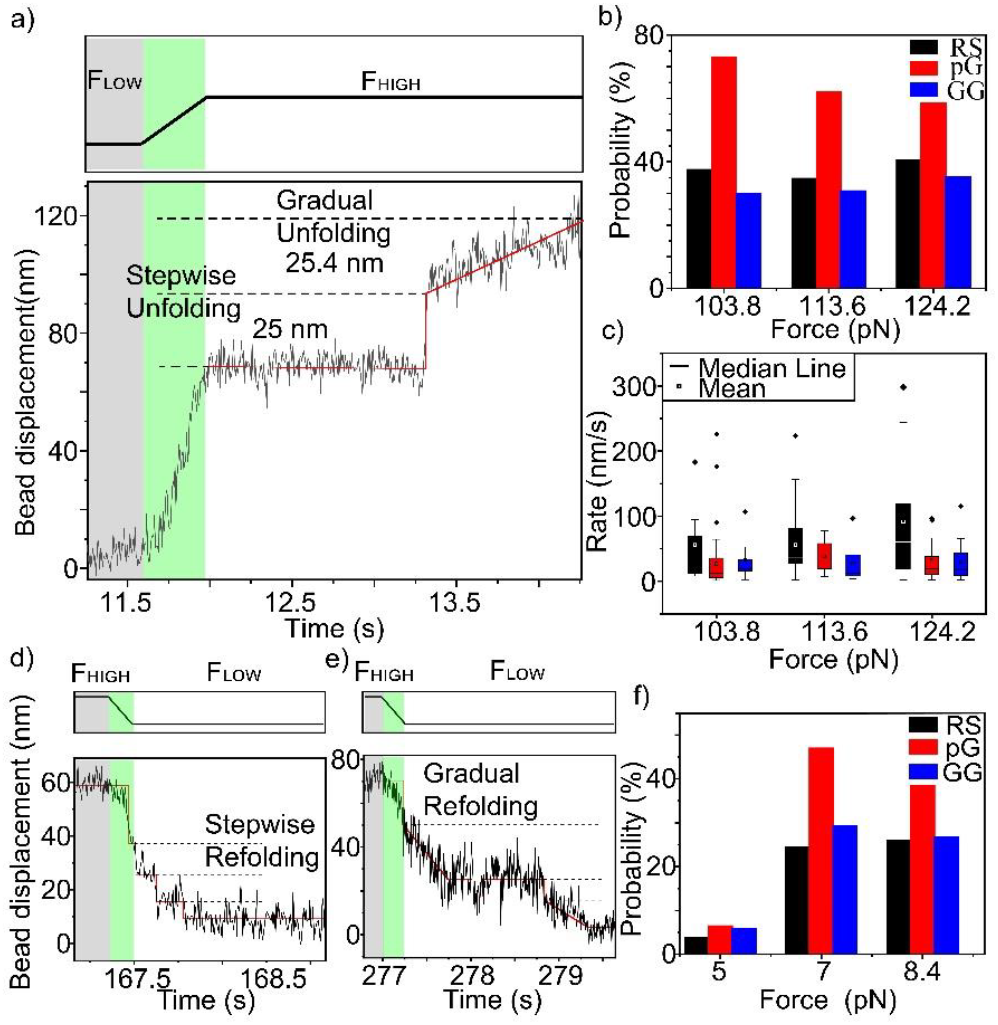
Creep analysis reveals linker dependence on cooperativity for all three models. a) A representative time trace of bead displacement along the z-axis during force clamp experiment. As we go from lower force (F_LOW_) to higher forces (F_HIGH_) for unfolding, we observe either step-like unfolding or very fast (about 0.5-2 seconds) gradual increase. The grey colored box highlights the region at low force and the green colored box represents the transition time to go from low force to high force. The red lines correspond to the step-fitting, which has significant goodness in fitting the step-like unfolding patterns but does not fit well for gradual increase. b) The probability of finding unfolding creep events at 103.8, 113.6 and 124.2 pN are shown using histogram plot for RS (in black), pG (in red) and GG (in blue). c) The rate of unfolding creep (creep length/creep lifetime) has been shown using a box plot at 103.8, 113.6 and 124.2 pN for RS (in black), pG (in red) and GG (in blue). The mean rate values are highlighted using a white box in the black box, black box in the red box and black box in the blue box for RS, pG and GG, respectively. Similarly, as we go from higher forces (F_HIGH_) to lower force (F_LOW_) for refolding, we observe either step-like refolding (d) or gradual decrease (e). The grey colored box highlights the region at high force and the green colored box represents the transition time to quench from high unfolding force to low refolding force. The red lines correspond to the step-fitting, which has significant goodness in fitting the step-like refolding patterns in (d) but does not fit well for gradual decrease in (e). f) The probability of finding refolding creep events at 5, 7 and 8.4 pN are shown using histogram plot for RS (in black), pG (in red) and GG (in blue).

Next, we measured the dwell-time and elongation lengths of creep events (Figure 3 a,d,e; SF 6-7) and estimated the corresponding creep rates (elongation length / dwell time). We measured the fastest mean unfolding creep-rate for the RS variant at all three clamping forces, followed by comparable creep rates of both pG and GG, respectively (Fig 3c).

Creep-rate is related to the cooperativity where faster creep-rate indicates more cooperativity and more synchronized unfolding transitions. This suggests interdependent structural rearrangements facilitated by unique chemical interactions between the linker and its domains in RS variant, potentially stabilizing intermediates and promoting efficient mechanical energy dissipation. Our in-silico findings also suggest the same (SF 8). Notably, cooperative behavior of RS variant is consistent across all forces, underscoring its distinct structural and chemical properties. Unfortunately, similar analysis of refolding creep-rate was limited due to significant overlap between force relaxation and refolding timescales during force quenching.

### In silico-studies

#### 1) Comparison among conformations

Next, we performed all-atom molecular dynamics (MD) simulations to understand the molecular mechanisms that determine the differences in mechano-stability and folding dynamics among variants. MD simulations were performed using the model structures generated from alphafold2 Colab (31, 32) (SF 9). From the simulations, we measured the radius of gyration (*R*_*g*_), interdomain bent angle (*θ*), interdomain torsional angle (*Φ*), and end-to-end distance (*L*_*end*_) (SF 10). We obtained comparable values of R_g_, θ, and L_end_ for RS and GG, indicating conformational similarity. In consistency, we obtained identical elution volumes for RS and GG variants (SF 11) and similar intra-domain interaction network for all three (SF 12). However, RS induces a more restricted *Φ* than the GG linker (Figure 4h). We infer that the bent geometry for RS and GG linker variants is attributed to shorter linker lengths. However, restrictions on the *Φ* for RS may be imposed due to persistent H-bonds between RS linker and domains (Figure 4l and Supplementary Table 4). pG, in contrast, displayed higher values for θ, Φ, L_end_, and R*g* (Figure 4d), suggesting an extended, more flexible, and distinct conformation compared to RS and GG (Figure 4f and 4h).

**Fig 4.**
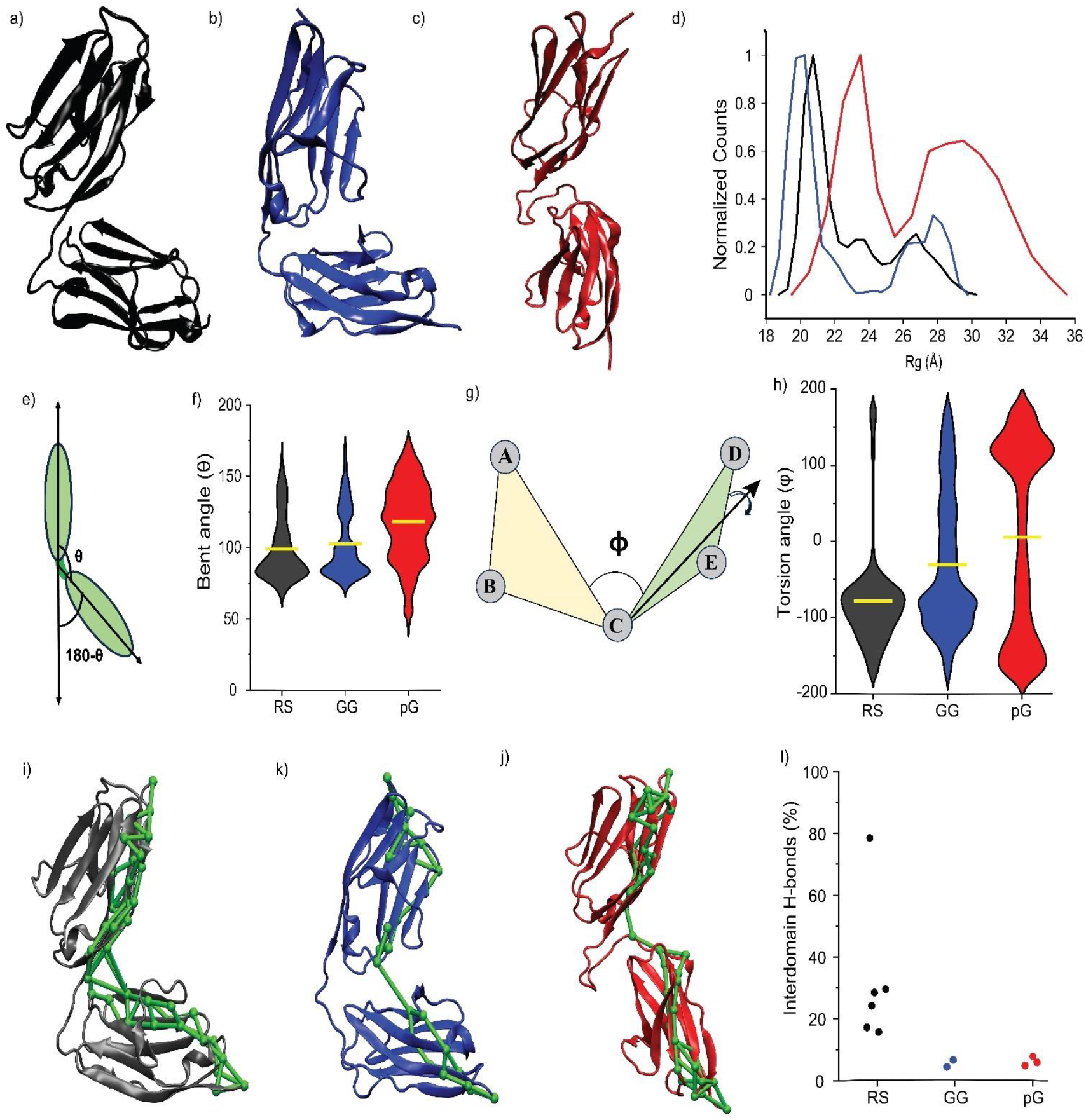
In-silico data shows difference in stability and flexibility among the three variants. (a), (b), and (c) cartoon representation ((RS, black), (GG, blue and pG, red)) of the most populated structure obtained from the Gaussian-accelerated molecular dynamics simulation. d) Radius of gyration plot, showing that RS and GG have similar compact conformation while pG has flexible and more extended conformation. e) Schematic representation of bent angle (θ) between domains and f) Violin plot for the bent angle, suggests RS and GG have similar distribution while pG has wider distribution. The yellow lines represent the mean value of the bent angle. g) Schematic representation of the torsion angle between planes and h) representing the distribution of torsion angle between two domains throughout the simulation, in all three proteins. The yellow line represents the mean value of torsion angle. RS has one rigid conformation while GG and pG have flexible conformation. (i), (j) and (k) All the sub-optimal paths (green) obtained from a dynamic network analysis of SMD pulling trajectory are overlaid on the ribbon structure (black, RS), (blue, GG) and (red, pG). Here, solid green tubes denote edges, and green spheres denote nodes. We repeated the SMD three times for each protein and took 100 frame trajectories for analysis (ignored the first 50 frames). We observed similar force propagation pathways in each run in all three systems. l) Scatter plot for the Interdomain H-bonds with their occupancy in all three proteins. This plot showing RS linker variant has five H-bonds with higher occupancy while the pG and GG linker variants also formed some transient interdomain H-bonds with smaller occupancy (<5 %). For, RS linker variant, we did not consider H-bonds those have occupancy less than 15 % of simulation time.

#### II) Comparisons of force-propagation pathways

Different degrees of restrictions in inter-domain orientations may steer the force propagations from one domain to next differently. To elucidate, we perform all-atom steered molecular dynamics simulations (SMD) within the experimental force-ramp range of 10^4^ µm/s and 5*10^3^ µm/s pulling velocities, respectively. We then estimated the force propagation paths using the Floyd–Warshall algorithm(33) (see Methods). To identify the key nodes responsible for force transmission, we analyzed the suboptimal paths of force propagation within a range of 20 nodes from the optimal path(34, 35). Our investigation unveiled a broader distribution of suboptimal paths in RS than pG and GG, indicating a more robust and effective dispersal of forces for RS in comparison to pG and GG (Figure 4i,k,j). Notably, the GG exhibited narrowest paths, highlighting its lowest mechanic stability and efficiency in disseminating forces and thus, featuring lowest mechanostability in experiments. The narrower distribution of paths transmits the external stimuli through a smaller number of nodes. Consequently, this reduces the dissipation of stimuli and renders the protein more susceptible to mechanical force.

### IDLs alter the protein longevity during repetitive stretch and release

The elastic properties of domains in mechanical proteins are associated with the storage and release of mechanical power under quasi-equilibrium conditions, as observed in constant force experiments(17). However, these proteins are often exposed to repetitive stress and release in cellular environments. While *constant* force-clamp measurements determine the thermodynamics and kinetics of probes along the pulling directions, the *repetitive force-release cycle* at a constant amplitude and frequency tests the non-stationary dynamics or the time-dependent mechanoresponsive behavior of a probe. We thus employed *oscillatory* force-clamp measurements where we set the mean-force (F_osc_) to 45 pN, amplitude (A) of 5 pN, and frequency (**ω**) of 0.25 Hz for a span of 60 minutes (data collected at every 5 mins interval and with a rate of about 200 FPS) using MT (Figure 5 a and b). We compare the time-dependent changes in power spectrum intensity across all variants. Since the domains of Titin are considered as the power-storage units in muscles, we wanted to estimate how linkers contribute in the power-retention ability of domains or in the stiffening of domains with time. Use of 45 pN as F_osc_ is not arbitrary. We wanted to monitor the time-dependent phenomena of the intact variants without completely unfolding, so decided to set the F_osc_ lower than 80 pN (as mentioned previously). To note, the estimated unfolding lifetime of I27 at 50 pN is ∼10^3^ s. Forces lower than 45 pN were avoided because the unfolding lifetime increases exponentially at lower tensile forces, and in the 20–30 pN range, the time required to observe significant deformation would exceed the experimental timeframe. Similarly, the choice of ω = 0.25 Hz balances the need to capture time-dependent phenomena while minimizing mechanical noise in the measurements, ensuring reliable data acquisition within the constraints of the experimental setup (SF13).

**Fig 5:**
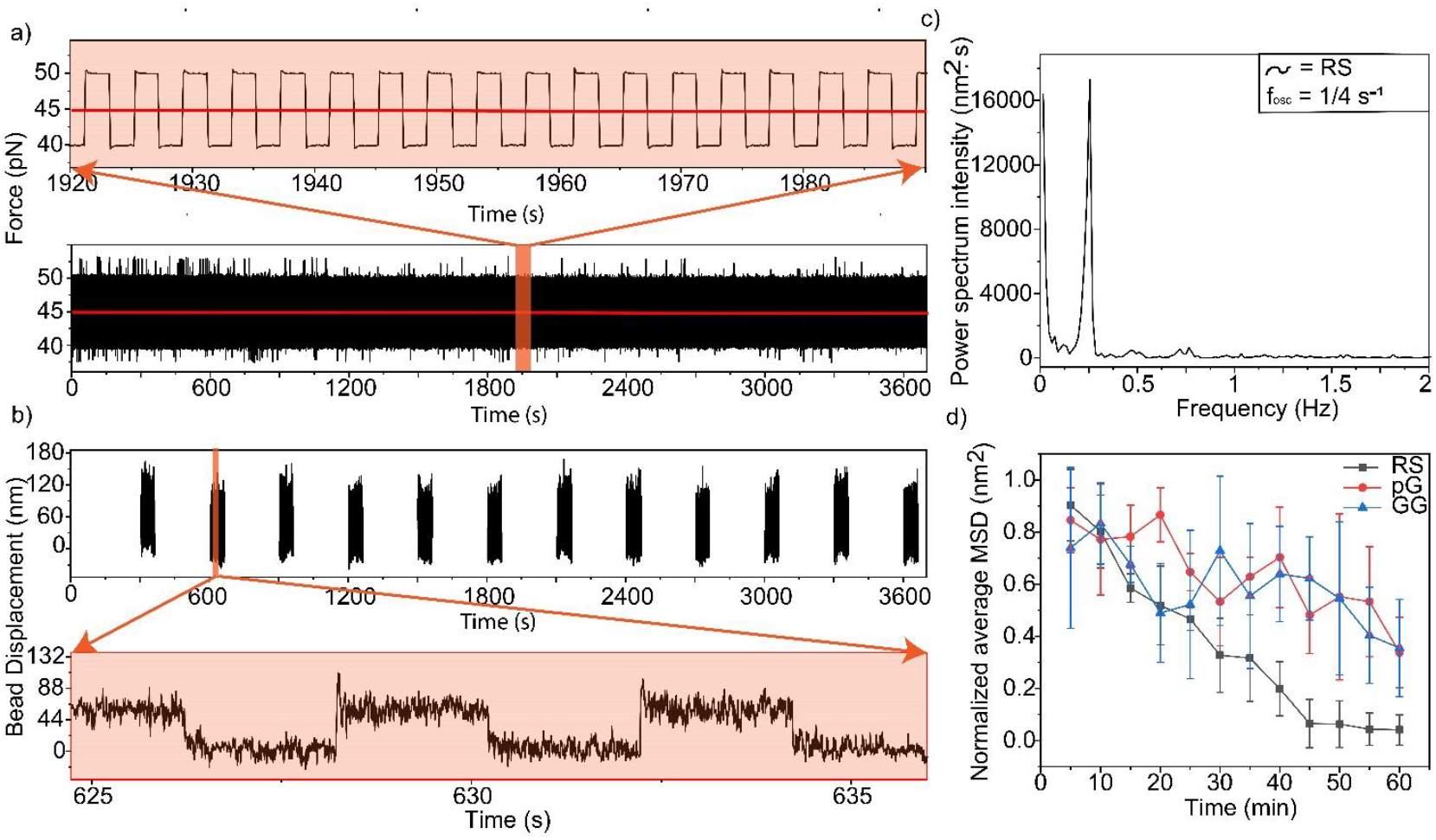
Repetitive stretch and release affect proteins folding and longevity. a) Representation of oscillation data collected for one minute after every five minutes from a I27 dimer with GGGSGGGG as IDL. It was repetitively stretched and released for one hour with a frequency of 0.25 Hz under a mean oscillatory force of 45 pN with 5 pN amplitude deviation. The data collection rate is about 200 Hz. a) A part of force-value data is highlighted in an orange box (about 70 seconds) and the zoomed-in version is shown. b) A part of bead position data is highlighted in an orange box (about 10 seconds) and the zoomed-in version is shown. c) Representation of power spectrum analysis of 60 seconds long extension versus time trace data. It is done for each of the one-minute time trace data which is collected after every five minutes and d) a normalized average intensity is plotted with respect to time for all three variants. The number of beads for this experiment was n=3 for pG, n=6 for GG, and n=3 for RS. The control has very short handles (PEG polymers), attached between bead and surface, which are not elastic element. This PEG stretch is entropic in nature and the signal output is stationary. As there is no decay in power output, we don’t observe any damping at all (SF 16).

From the force-spectrum (extension-time traces), we measured the power spectral density for a 60 s trace collected at every 5-min interval and monitored the change in energy (Area under the power spectrum) with time (see Methods, SF14, Figure 5c). We observed a decay in energy with time for all three variants, a phenomenon commonly known as damping in material science (SF15, Fig 5d). For RS, the intensity drop is significantly faster, like overdamped condition, than other variants (Fig 5d). pG and GG variants behaved underdamped. For quantitative understanding, we assume all three variants as underdamped harmonic oscillator and fit to the mechanical energy decay equation: E(t)=E_0_e^−D.t^, where D (damping-coefficient per unit mass) = 2.ζ.ω_0_ (ζrepresents damping-ratio and ω_0_ is the natural frequency of oscillation). We measured the highest D for RS (0.061 ± 0.005 s^-1^) followed by pG (0.012 ± 0.002 s^-1^) and GG (0.01 ± 0.003 s^-1^) (Fig 5e). RS variant is mechanically most stable, and thus stiffest, as obtained from constant force measurements. The increased stiffness raises the natural frequency of the spring, which can lead to quicker responses. Meanwhile, the higher damping rapidly dissipates energy, reducing overshoot and oscillations. This combination leads to a more resilient and responsive system, especially useful in environments where both high load-bearing capacity and rapid vibration control are essential. On the contrary, GG induces lower damping coefficient and lower stiffness. While lower stiffness may allow the GG variant to respond to smaller forces with greater displacement, lower damping facilitates less energy dissipation as heat during oscillations, allowing the system to store and release energy more efficiently. Such a spring with lower stiffness and damping is ideal for applications where sensitivity, smooth operation, and efficient energy utilization are prioritized over rapid settling or high-load support.

## Discussion

The intricate interplay between interdomain linkers (IDLs) and protein domains in multidomain mechanical proteins, such as titin, has long been overshadowed by the focus on individual domains. This study reveals that IDLs are not passive connectors but dynamic modulators of mechanical stability, energy retention, and fatigue resistance, offering critical insights into how mechanical proteins adapt to diverse force environments. By engineering titin-derived I27 dimers with distinct IDLs (rigid RS, short GG, and flexible pG), we demonstrate that linker properties dictate conformational dynamics under both constant and oscillatory forces, with profound implications for biological function and synthetic protein design. Our findings bridge nanoscale protein mechanics to tissue-level function, while underscoring IDLs as overlooked yet pivotal players in muscle physiology and disease.

Titin’s modular architecture exemplifies a universal trade-off in mechanobiome proteins: rigid IDLs (e.g., RS) enhance mechanical stability but compromise energy retention under cyclic loading, while flexible linkers (e.g., pG) prioritize power conservation at the cost of rapid fatigue. During muscle contraction, RS-like rigidity allows titin to withstand peak stresses (>4 pN) through cooperative, stepwise unfolding transitions stabilized by H-bonds and restricted torsional angles. Conversely, pG-like flexibility enables creep-like deformations—gradual, non-cooperative transitions that dissipate energy slowly, prolonging titin’s role as a power reservoir. This duality mirrors muscle’s need for both explosive force (rigid phases) and endurance (flexible phases), with IDLs acting as tunable “dials” to optimize performance across physiological demands.

Notably, disease-associated mutations disproportionately target IDLs over domains (e.g., in titin, spectrin, and cadherins), disrupting their mechanical filtering capacity. For instance, spectrin linker mutations destabilize erythrocyte membranes(36), causing hemolytic anemia, while titin IDL variants correlate with cardiomyopathies(37). Our work rationalizes these pathologies: mutations altering linker flexibility or interactions likely perturb the balance between energy retention (critical for sustained contraction) and fatigue resistance (vital for repetitive motion).

A key discovery is the prevalence of creep-like transitions in titin—a phenomenon classically associated with materials science(38). These second-order transitions, dominant in pG-linked dimers, involve gradual, reversible shifts between metastable microstates rather than binary folding/unfolding. Creep enables titin to absorb mechanical energy without irreversible damage, akin to viscoelastic polymers. Flexible IDLs promote this behavior, as seen in pG’s higher creep percentage and superior power retention under oscillatory forces. We propose that creep-like dynamics consume less energy than cooperative transitions, allowing titin to buffer stochastic forces during muscle activity. Conversely, RS’s rigid, cooperative transitions prioritize rapid energy dissipation, ideal for load-bearing but prone to fatigue—a trade-off mirrored in synthetic dampers.

Molecular dynamics simulations reveal how IDLs govern force transmission. The RS linker’s hydrogen-bonded interactions with domains distribute stress across broad, redundant pathways, mitigating localized damage—a design principle akin to carbon-fiber composites. This network-level robustness underpins RS’s superior stability under constant force (SF 8). In contrast, GG’s narrow force pathways concentrate stress, hastening failure. Strikingly, RS’s restricted torsional angles and interdomain crosstalk also accelerate refolding, positioning IDLs as allosteric regulators that couple mechanical input to conformational output.

Under oscillatory forces, however, RS’s overdamped behavior—rapid energy loss via internal friction—contrasts with pG’s underdamped resilience. This dichotomy mirrors engineered systems: stiff alloys resist deformation but fatigue faster than flexible polymers. In muscle, such linker-dependent damping may allow titin to dynamically tune stiffness during contraction-relaxation cycles(39), ensuring efficient energy transfer while avoiding catastrophic failure.

Our findings challenge the myosin-centric view of muscle contraction by positioning titin as a co-driver of mechanical power(17). Flexible IDLs enable titin to match myosin’s energy output(17) during repetitive motion, while rigid linkers provide passive stiffness during isometric holds. This synergy is disrupted in diseases like cardiomyopathies, where IDL mutations may impair titin’s ability to switch between energy-storage and stability modes. Furthermore, the heightened fatigue susceptibility of rigid linkers under oscillatory forces (e.g., RS’s rapid damping) underscores why aging or oxidative damage—which stiffen proteins—predispose tissues to mechanical failure.

The IDL-mediated trade-off between stability and adaptability offers a blueprint for synthetic protein design. RS-like linkers could stabilize biomaterials in load-bearing implants, while pG-like flexibility might enhance energy efficiency in soft robotics. Additionally, targeting IDLs with pharmacological chaperones could rescue pathogenic mutations in diseases like hereditary elliptocytosis(40). Future work must explore how post-translational modifications (e.g., phosphorylation) dynamically tune linker mechanics in vivo and how multi-domain chains integrate diverse IDLs for emergent functions.

## Conclusion

This study redefines IDLs as central architects of protein mechanics, orchestrating titin’s dual role as a shock absorber and power generator in muscle. Their ability to balance cooperative rigidity with creep-like plasticity reflects an evolutionary optimization for diverse mechanical landscapes. By bridging protein folding theory, materials science, and physiology, we illuminate how subtle variations in linker chemistry enable life’s dynamic mechanics—from the beat of a heart to the leap of an athlete. These insights not only advance our understanding of muscle disease but also inspire a new generation of biomaterials that emulate nature’s mechano-adaptive genius.

## Material and methods

### Cloning, protein expression and purification

Two repetitive I27 units with RS, GG and pG as interdomain linkers are cloned between Nde1 and Xho1 restriction sites in pET22b vector.At the C-terminus of every construct is a 6xHis-tag and a sort-tag (LPETGSS). His-tag was required for affinity-based protein purification, while Sort-tag was introduced for covalent attachment to the surface. N-terminus contains an Avitag for in-vitro biotinylation of protein. The RIPL Escherichia coli expression strain has been used to express proteins for the studies. Cultures were grown in LB at 37°C until they achieved an OD_600nm_ value of 0.6. Protein expression was induced by adding 1mM IPTG at 37°C for 4-5 hours. The cells were harvested by centrifugation at 6000 rpm and 4°C.The cell pellets were resuspended in lysis buffer (50 mM potassium phosphate, 300 mM KCl, pH 7.5) in presence of protease inhibitor cocktail and lysed by sonication at 4°C. The supernatant upon centrifugation (12000 rpm for 30 minutes at 4°C) was purified using Ni-NTA chromatography. The resin was washed with a phosphate buffer followed by 20 mM imidazole (prepared in phosphate buffer). All the protein constructs with Avitag at the N-terminus and His6 tag at C-terminus were eluted using 200 mM and 500 mM imidazole (in phosphate buffer). SDS-Page was used to validate the protein’s purity, and the protein bands were found to be at appropriate size. Finally, the proteins were further purified using fast-performance liquid-phase chromatography (FPLC) in 25mM HEPES, 50mM NaCl, 25mM KCl buffer at pH 7.5. Western blotting with conjugated streptavidin horseradish peroxidase (GE) was used to confirm the biotinylation of the protein.

### Size Exclusion Chromatography (SEC)

The Superdex 200 (Cytiva) column coupled to an AKTA Pure system was used to perform analytical size-exclusion chromatography (SEC). The size exclusion chromatogram reveals that (I27-RS-I27) and (I27-GG-I27) have similar molecular sizes, whereas (I27-pG-I27) exhibits a slightly larger size.

### Magnetic Tweezers Setup

Our lab built magnetic tweezer setup is constructed on top of an inverted microscope (Olympus IX73).The tweezer setup comprises a pair of N52 grade neodymium permanent magnet, high precision voice-coil actuator (VCA), a high-speed camera, an inverted microscope, and a Python-based algorithm for image analysis and data acquisition. VCA has a highly accurate feedback mechanism with a resolution of 0.01 mm to exert a precise force on the protein under observation. We use VCA as a linear actuator to move the magnet position and exert force on the superparamagnetic beads (2.8 µm diameter, Fe_3_O_4_) attached to the surface via the protein to be probed. The particle moves up and down under the influence of magnetic force that approaches the fluid chamber from top. The temporal Z-displacement of the magnetic particle features the unfolding and refolding patterns of the probe protein under varying forces.The CMOS camera records bead movement in the z-direction under force, and an algorithm based on Python is used to process the data in real time. The list of the equipment used in the setup are mentioned below.

1. BEI Kimco Voice Coil Actuator (VCA) Developer’s Kit (DK-LAS13-18-000A-P01-3E), which includes an Ingenia Motion Controller (PLU-1/5-48C)
2. Piezo-scanner (P-725.xDD PIFOC) and controller (PI-E-709).
3. Inverted Optical Microscope with 100x objective (Olympus IX73)
4. High-Speed CMOS camera (Ximea xiQ USB 3.0 SuperSpeed)
5. Pair of Neodymium Permanent Magnets

### Surface modification protocol for Single-molecule force spectroscopy

The glass coverslips were subjected to air plasma activation and then treated with piranha solution (v/v 75% H2SO4, 25% H2O2) for 45 min, followed by a thorough rinse with deionized water. Silanization of coverslips with 1% APTES (3-Aminopropyltriethoxy silane, v/v) takes place in 95% acetone for 45 min. Coverslips were washed with acetone by sonication, followed by rinsing with water, and repeating the process with acetone. Subsequently, coverslips were placed in a vacuum oven at 100°C for 1 hour. Amine-exposed coverslips were treated with NHS-PEG2-Maleimide in basic buffer (100 mM NaHCO3, 600 mM K2SO4, pH 8.3) for 3 hours. Then, coverslips were washed with deionized water to remove the unreacted substrate. The PEGylated surfaces were thereafter subjected to a 7-hour room temperature (RT) incubation with 100µM polyglycine peptide, GGGGC, in order to facilitate the cysteine-maleimide reaction. Sortase-based covalent attachment uses polyglycine on coverslip as a nucleophile. Coverslips were rinsed with water and kept in a vacuum desiccator prior to protein attachment.

Polyglycine(GGGGC) exposed coverslips were incubated with protein of interest (POI), I27 here, and Sortase in a 4:5 molar ratio to facilitate the formation of a thioester intermediate in 2mM Ca2+, 25mM HEPES, 50mM NaCl, 25mM KCl buffer at pH 7.4 at room temperature (RT) for 1.5 hours. Unreacted Sortase and POI were removed from the surface by washing it with a buffer.Sortagging reaction covalently attaches POI (I27)_2_ to the cover slip. Protein linked to the coverslip was incubated with a mixture of 10 um BirA (biotin ligase), 10mM ATP, 10mM MgCl_2_, 50mM Bicine buffer (pH 8.3), and 500uM Biotin for 4 hr at RT. Avitag is present at the N-terminus of the protein, and this reaction performs the in-vitro biotinylation of Avitagged protein. Magnetic beads (Dynabeads M270-Amine) modified with Streptavidin were incubated with biotinylated protein for 20 min.The last step is the attachment of non-magnetic polystyrene beads to the coverslip as a reference to correct the spatial drift during the experiment. 5ul of Non-magnetic Amino Polystyrene Beads (2.67um size) diluted with 95 ul buffer were incubated on the surface for 20 min to allow non-specific adsorption. After 20 min, coverslips were washed with a buffer. Cover slips are ready for use in the Magnetic tweezers experiment.

### Magnetic Tweezer Experiments

The magnetic tweezers allow us to use a magnetic field to exert controlled forces on magnetic beads attached to biological molecules. These beads are typically microscopic (radius of 2.7 microns) and can be manipulated with high precision. These superparamagnetic beads are made of a magnetic material (e.g., iron oxide) that are attached to biological molecules, here I27 dimer. The beads are usually coated with a surface layer (e.g., streptavidin) to facilitate binding avitag on the proteins N-terminal. The other end, C-terminal is anchored to a surface (e.g., a glass slide) using a sort-tagging chemistry. Neodymium grade permanent magnets create a magnetic field. The field’s strength and direction can be controlled to manipulate the position and orientation of the magnetic beads. A stage that can be moved with 0.01 mm precision, often equipped with piezoelectric actuators, with a computer control system that adjusts the VCA position to manipulate the field strength. A high-resolution microscope with a 100x objective lens is used to observe the movement of the magnetic beads and the attached molecules. Often, video microscopy is employed to capture real-time data. However, here we apply a z-stacking program for every bead we want to observe and use an image processing algorithm to collect real-time data.

The buffer conditions were kept the same throughout the experiment. It was 25mM HEPES, 50mM NaCl, 25mM KCl buffer at pH 7.5. Data was recorded at 200 Hz. We employed two types of designs 1. Constant force clamp 2. Oscillatory force clamp

Time traces of bond survival were collected at each clamping force to calculate bond survival probabilities. The probability of the states at respective force-clamps was estimated from the dwell time (Dwell-Time Distribution Analysis of Polyprotein Unfolding Using Force-Clamp Spectroscopy) of each state, normalized to the total clamping time. Force clamp experiments were performed at three clamping forces (103.8 pN, 113.6 pN and 124.2 pN) for unfolding. Similarly, refolding was performed at three different force values (5.0, 7.0, 8.3 pN).

The I27 dimer was subjected to mean oscillatory force perturbations at 45 pN with 5 pN amplitude (between 40 and 50 pN) for 60 min at 0.25 Hz frequency using MT. The power spectral intensity was measured from the time vs extension trace using a periodogram program. Later on, a Lorentzian fitting was performed, to fit the data across all possible frequencies and give their intensity. A specific magnetic bead is verified from i27 fingerprint before starting the force-oscillation experiment. For the control experiment we have taken a magnetic bead and attached that directly to the surface without a protein of interest in between.

### Data Analysis

After data acquisition, drift correction was carried out with the help of reference beads. Step fitting technique was used to further process the drift-corrected data. Auto-stepfinder, an open-source resource, a GUI based MATLAB programme was used for step fitting analysis(28). The difference in time between the application of the clamping force and the changes in the native or denatured state of the protein was used to compute the lifetime of each state. Fitting the survival probability with exponential decay functions yields the mean lifetime of the state at each clamping force and this analysis was performed using OriginPro software. The survival probabilities are best fit to mono-exponential decay in case of pG and GG whereas bi-exponential model fits for RS at low force values. The details of the fit are given in SI table (2 & 3)

### Gaussian-accelerated molecular dynamics simulation

System for the RS, GG, and pG linker variants of I27 were prepared using QwikMD(41) plugin within VMD(42). The structure was aligned to the longest z-axis and solvated using TIP3 water in a box maintaining the distance of 15 Å from the edges of the box. The box dimensions were of 61.03 Å × 123.15 Å × 62.55 Å in the X, Y, and Z directions, placed at coordinates (−29.89, -63.04, -31.69) and (31.14, 60.11, 30.86) as the minimum and maximum coordinates. Following solvation, Na+ and Cl− ions were randomly placed in the water box to achieve a concentration of 150 mM, replacing water molecules. The simulations were carried out with NAMD(43, 44) version 2.12 employing the CHARMM36 force field(44). Model structures for all three variants were generated using alphaFold2 Colab, and the alphaFold-generated model served as the starting structure for our simulations. The setup and simulation protocol were identical for all three systems.

The system underwent minimization for 5000 steps using a conjugate gradient minimization algorithm. Subsequently, the system’s temperature was slowly raised from 60 K to 300 K at a rate of 1K every 600 steps over 0.29 ns, with backbone atoms restrained. This was followed by a 2 ns equilibration period with the backbone atoms restrained. The equilibrated system was used for Gaussian accelerated molecular dynamics simulations(34, 44, 45), which included 5 ns of classical molecular dynamics followed by 20 ns of equilibration during which boost potentials were collected. Two independent repeats of 120 ns Gaussian accelerated simulations were then performed with fixed boost parameters. All simulations were carried out in dual-boost mode with the boost potential set to Vmax (maximum potential energy). Throughout these steps, the system’s pressure was maintained at 1 atm using the Nose–Hoover piston(46, 47). Long-range interactions were treated using the particle-mesh Ewald (PME) method(47), and the r-RESPA scheme was employed to update short-range van der Waals interactions every step and long-range electrostatic interactions every two steps. The trajectories were concatenated using the software Catdcd 4.0, and end to end distance was calculated between the alpha carbon atoms (Cα) of first and last amino acids using the Euclidean distance formula in VMD. The radius of gyration defines the root mean square distance of all atoms or amino acids in a molecule from the center of mass were calculated using a script in VMD. The inter-domain angle refers to the angle between the vectors projecting from the 1^st^ to 89^th^ residue in each domain throughout the simulation. We analyzed hydrogen bonds using a donor-acceptor distance cutoff of 3Å and an angle cutoff of 20 degrees in VMD.

### Steered molecular dynamics simulation

The steered molecular dynamics (SMD) simulations were carried out using the most populated conformation (Figure .3 (a), (b) and (c)) obtained from the Gaussian accelerated MD simulation. The setup and simulation protocols were kept identical across all three systems. Initially, the structures were aligned along the Z-axis. Subsequently, the system underwent solvation by immersing it in a water box with a 15Å buffer around it, extending 100Å in the pulling direction. The dimensions of the water box 83.93, 83.93, 298.38 Å in X, Y, and Z directions placed at (−43.24, -43.40, -48.42) and (40.70, 40.54, 249.96) as the minimum and maximum coordinates and TIP3P water used for the solvation. The Na^+^ and Cl^−^ ions corresponding to a concentration of 150 mM were then placed randomly in the water box by replacing the water molecules. The Cα atom at the C-terminus of proteins was held fixed, while the Cα atom at the N-terminus was pulled at constant velocities of 0.01nm/ns and 0.005nm/ns with a spring stiffness of 7 kcal mol^-1^A^-2^. SMD was executed by applying harmonic restraints to the position of LEU180 for RS and GG proteins, LEU186 for pG, along with a secondary restraint on LEU1. We repeated the SMD three times for each protein and took a 100 frames trajectory after 50 frames for calculating the force propagation pathway. Force propagation pathways were calculated using the Floyed-Warshall algorithm. This algorithm identifies the shortest paths between interacting nodes in a protein structure based on a distance matrix where each entry represents the minimum distance between the nodes, i.e., the strength of an edge. The algorithm iteratively updates the distance matrix by considering each node as an intermediate point. For each pair of nodes (i, j) the algorithm checks whether the path from i to j through an intermediate node K is shorter than the direct path from i to j. If so, it updates the matrix to reflect the shorter path. This process is repeated for all possible pairs of nodes and all possible intermediate nodes. After considering all nodes as intermediates, the distance matrix will contain the shortest paths between all pairs of nodes.

### Dynamic network analysis

The Dynamic Network Analysis was conducted using the network view(34) plugin in VMD along with associated scripts. In the network, nodes were created based on α-carbon atom of each residue and edges drawn between them if the distance between two nodes was within a cut-off distance of 4.5 Å for at least 75% of the trajectory. The weight of the edges was determined using the correlation matrix generated by the program “carma”(48). Edge strength between nodes was calculated using pairwise correlation (C_ij_), reflecting the probability of information transfer between them with the equation w_ij_ = −log (|c_ij_|). Notably, neighboring α-carbon atoms, same residues and hydrogens were excluded from consideration because they give non-trivial paths. The resulting network was then partitioned into sub-networks or communities using the Girvan–Newman algorithm(49). Then we computed the degree of each residue in all three proteins. The degree represents the extent of interaction of each residue with its neighboring residues or the number of nodes connected to a particular node. The data indicate that most of the residues in RS, GG, and pG linker variants exhibit a similar extent of degrees (Figure 7). This indicating that in all three systems, linkers do not change the interaction patterns within the domains.

### Force calibration using the DNA B-S overstretching(20, 29)

A DNA construct composed of 605 bp DNA segment is tethered between a coverslip and a superparamagnetic bead. The B−S transition is observed during the force ramp (n = 13) marking the 65 pN point. Histogram of magnet position values where the B−S transition occurred was plotted and the peak position was measured from gaussian fitting, which showed magnet distance of 4.31 ± 0.02 mm(20). Bin width was calculated using Freedman-Diaconis rule, 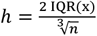, where h is the bin width, and n is the total number of data in x dataset (n=13)(20).

## Acknowledgements

We thank Ojas Singh for writing codes for Magnetic Tweezers. We thank Vishavdeep Vashisht for his help in building and standardizing Magnetic tweezers instrument. We thank Shaliza Kuttassari for her help in force-calibration. We thank Dr. Anil K.D. for his suggestions. This study was supported by DST Core Research Grant.

## Notes

### Competing Interest Statement

The authors have declared no competing interest.

