## Supplementary information for "Titin as a mechanical damper: Balancing Stability and longevity through inter-domain linker design"

[#]equal author contribution,

*Corresponding author

**SI Table**

Supplementary Table 1:  Propensity of disease mutations at linker regions of Titin.

Supplementary Table 2: Comparison of intrinsic folding and unfolding rates and transition states distance

Supplementary Table 3: Force dependent unfolding lifetime data and the goodness of fitting for each I27 doublet linker variant.

Supplementary Table 4: Force dependent lifetime data for refolding for I27 doublet linker variant.

Supplementary Table 5: Interdomain H-bonds and their occupancy in RS, GG, pG variants of I27.

**SI Table 1. Propensity of disease mutations at linker regions of Titin.**

| Disease | Mutation | |
|---|---|---|
| | Domain | Linker |
| Cardiomyopathy (CM) | 1.2% | 0.35% |
| Centronuclear myopathy (CNM) | - | 30% |
| Muscle disease (MD) | 5.68% | 2% |
| Hypertrophic cardiomyopathy (HCM) | 0.9% | 8.17% |

**SI Table 2 : Comparison of intrinsic folding and unfolding rates and transition states distance.** This table is made from the force-clamp data coming from three linker models to quantify differences in kinetics among variants. Values are calculated from $\ln \mathbf{k}_{f} = \ln \mathbf{k}_{0} + \mathbf{F} * (\mathbf{X}_{\beta} / \mathbf{k}_{B}.\boldsymbol{T})$. From the transition state distance from folded and unfolded state, $X_\beta$ of unfolding and folding gives estimation of force tolerance.

| Protein Variants | Fold-rate $k_0^f$ (s$^{-1}$) | Force-tolerance of folding, $x_\beta^f$ (nm) | Unfold-rate $k_0^u$ (s$^{-1}$) | Force-tolerance of unfolding, $x_\beta^u$ (nm) |
|---|---|---|---|---|
| RS | 6.29 ± 2.66 | -2.00 ± 0.19 | 0.003 ± 0.000 | 0.12 ± 0.0 |
| GG | 1.06 ± 0.75 | -1.24 ± 0.42 | 0.053 ± 0.018 | 0.04 ± 0.1 |
| pG | 1.12 ± 0.62 | -1.39 ± 0.33 | 0.038 ± 0.001 | 0.04 ± 0.1 |

**SI Table 3: Force dependent unfolding lifetime data and the goodness of fitting for each I27 doublet linker variant.**

**Table 3a. Force dependent lifetime of RS variant, extracted from exponential fit to the survival probability.**

| **Clamping force(pN)** | **lifetime(s)** | **Amplitude** | **Adj. $R^2$** | **Mean lifetime(s)** |
|---|---|---|---|---|
| 103.8 | T1 = (33.75±8.12)<br><br>T2 = (4.66±0.61) | A1 = 0.33<br><br>A2 = 0.67 | 0.988 | 14.25 |
| 113.6 | T1 = (23.13±12.51)<br><br>T2 = (4.77±1.15) | A1 = 0.30<br><br>A2 = 0.70 | 0.977 | 10.27 |
| 124.2 | T = (7.7±0.14) | A = 1.00 | 0.987 | 7.70 |

**Table 3b: Force dependent lifetime of pG variant, extracted from exponential fit to the survival probability.**

| **Clamping force(pN)** | **lifetime(s)** | **Amplitude** | **Adj. $R^2$** |
|---|---|---|---|
| 103.8 | T = (8.06±0.1) | A = 1.0 | 0.99 |
| 113.6 | T = (7.25±0.12) | A = 1.0 | 0.98 |
| 124.2 | T = (6.4±0.1) | A = 1.0 | 0.99 |

**Table 3c. Force dependent lifetime of GG variant, extracted from exponential fit to the survival probability.**

| Clamping force(pN) | lifetime(s) | Amplitude | Adj. $R^2$ |
|---|---|---|---|
| 103.8 | T = (5.90±0.13) | A = 1.0 | 0.98 |
| 113.6 | T = (5.60±0.12) | A = 1.0 | 0.98 |
| 124.2 | T = (4.72±0.1) | A = 1.0 | 0.98 |

**SI Table 4. Force dependent lifetime data for refolding for I27 doublet linker variant.**

| Clamping force (pN) | lifetime(s) | | |
|---|---|---|---|
| | RS | pG | GG |
| 5.0 | 2.04±0.02 | 5.21±0.06 | 4.66±0.1 |
| 7.0 | 4.3±0.06 | 8.2±0.14 | 6.44±0.12 |
| 8.4 | 8.94±0.07 | 17±0.43 | 13.57±0.33 |

**SI Table 5. Interdomain H-bonds and their occupancy in RS, GG, pG variants of I27.**

| RS | | pG | | GG | |
|---|---|---|---|---|---|
| **H- bonds** | **% Occupancy** | **H- bonds** | **% Occupancy** | **H- bonds** | **% Occupancy** |
| R90-E118 | 78.5 | L65-G95 | 7.8 | N168-Q64 | 6.7 |
| E88-D120 | 28.5 | G66-G96 | 5.9 | K170-D46 | 4.5 |
| K87-E142 | 24.2 | K85-D124 | 4.9 | | |
| S122-D88 | 29.6 | | | | |
| K85-D120 | 17.2 | | | | |
| K85-S122 | 15.7 | | | | |

**Supplementary Figures**

Supplementary Figure 1: Measurement of Chemical and thermal stability.

Supplementary Figure 2: Lab-built Magnetic tweezer schematics and picture.

Supplementary Figure 3: Contour length distribution from constant force-clamp study

Supplementary Figure 4: Exponential decay fitting of survival plots from native state.

Supplementary Figure 5: Exponential decay fitting of survival plots from unfolded state.

Supplementary Figure 6: Rate of creep like transition for all three variants

Supplementary Figure 7: Creep length distribution for all three variants

Supplementary Figure 8: Graphical representation of the contacts or interactions between amino acid residues in RS, pG, and GG linker variants shown by contact map

Supplementary Figure 9: RMSD plot for RS (black), GG (blue), and pG (red) variants: A comparison of structural dynamics

Supplementary Figure 10: Conformational dynamics of RS (black), GG (blue), and pG (red) variants: Insight from end-to-end analysis

Supplementary Figure 11: Comparative size-exclusion chromatogram (SEC) of the I27 doublet variants and CD spectra

Supplementary Figure 12: Comparison of residue interactions (degree analysis) in RS, Pg, GG linker variants with respect to RS variant

Supplementary Figure 13: Plot of 60-second traces recorded at 200 Hz frame per seconds after every 300 seconds while the beads are subjected to oscillatory force-pulse of 0.25 Hz.

Supplementary Figure 14: Power spectrum analysis example

Supplementary Figure 15: Decrease in power with time for all three variants

Supplementary Figure 16: Example of data, power spectrum analysis, and SNR plot from Non-specific magnetic bead.

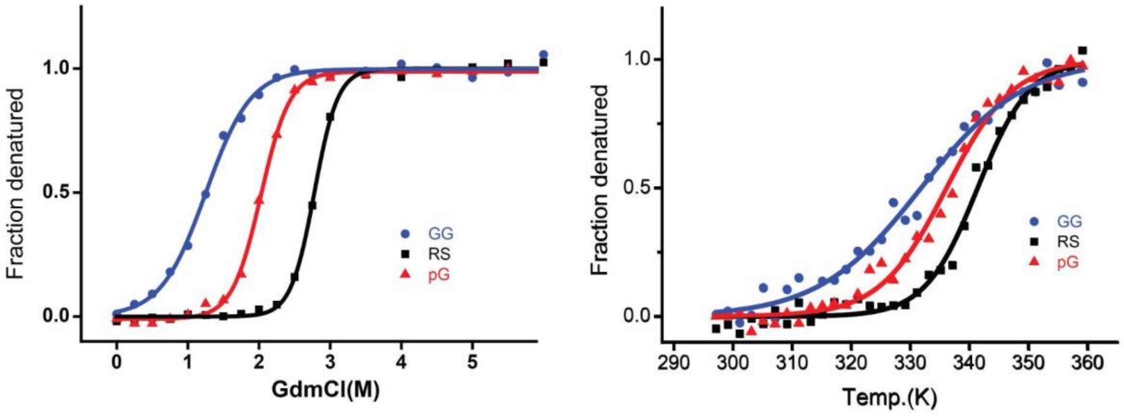


**SF 1: Measurement of Chemical and thermal stability:** Chemical and thermal unfolding of I27 doublet variants in presence of RS, GG, and pG as seen by intrinsic tryptophan fluorescence and far-UV CD respectively. To estimate unfolded populations, the data are normalized. The Boltzmann fits to the data are shown as solid lines.

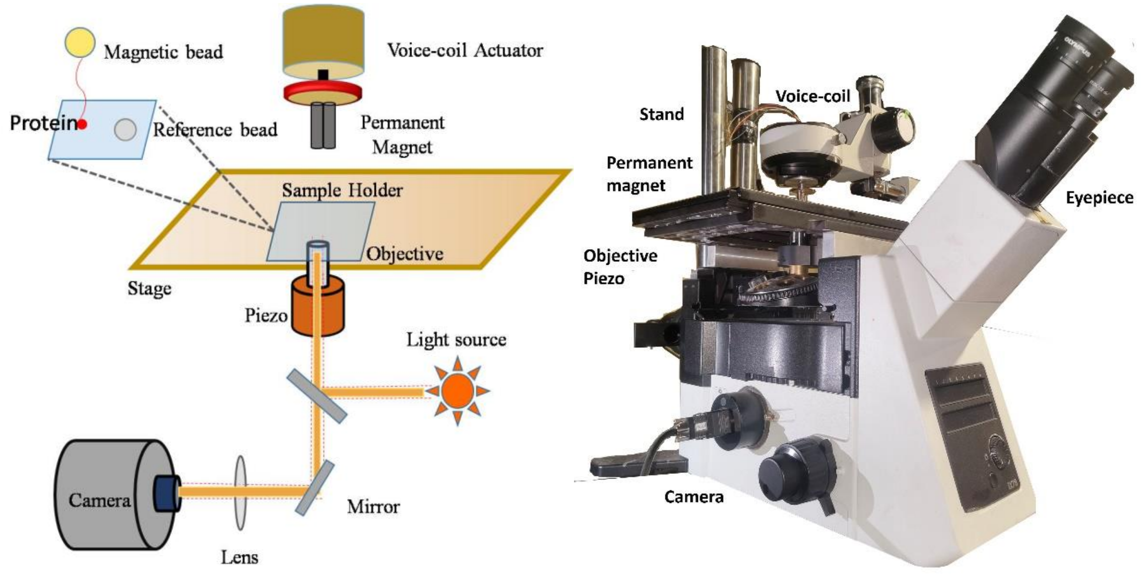


**SF 2: Lab-built Magnetic tweezer schematics and picture:** Representative diagram of magnetic tweezer setup is shown. Cartoon-diagram of the schematics of a magnetic tweezer components and light-path is highlighted. The actual picture of magnetic tweezers is also attached here.

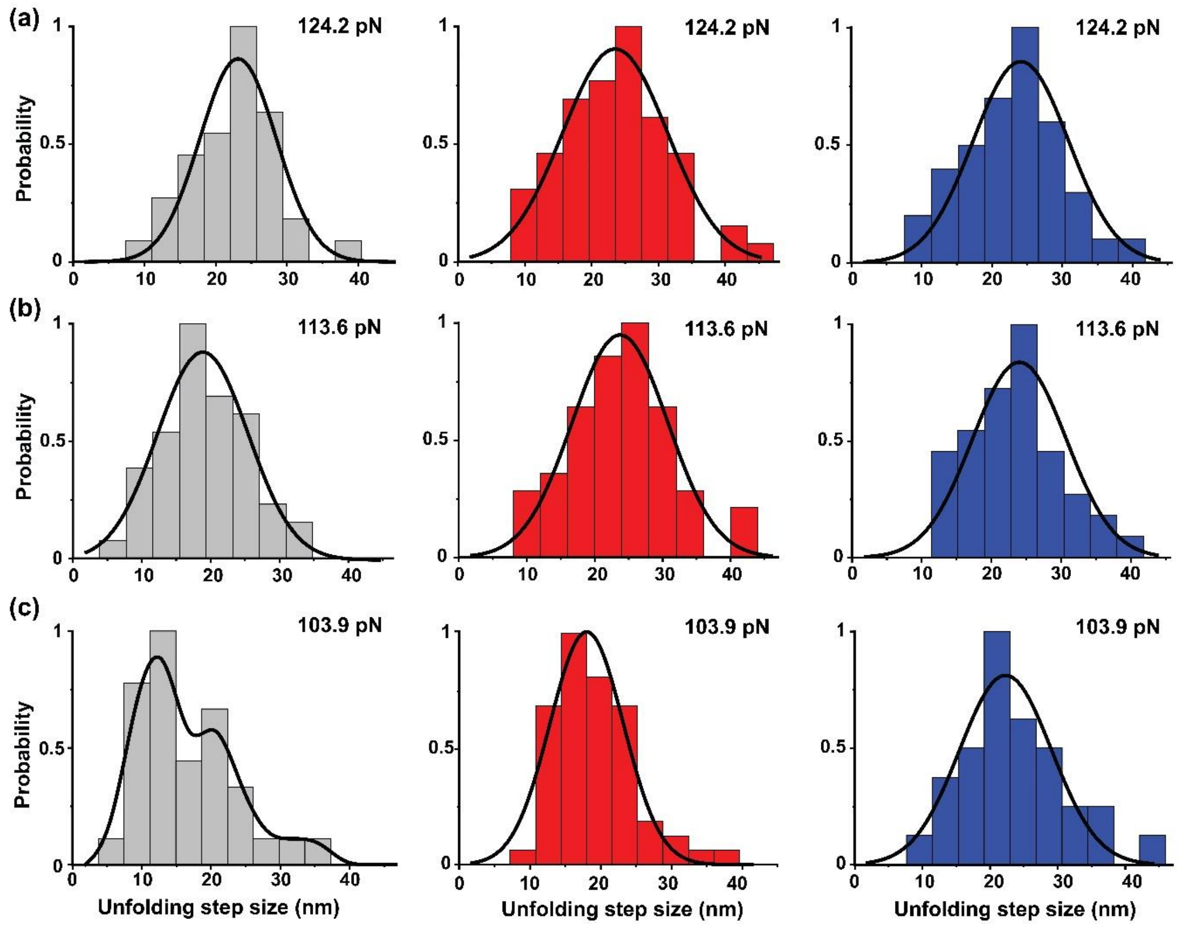


**SF 3: Contour length distribution from constant force-clamp study.** Distributions of unfolding length at 103.8, 113.6 and 124.2 pN clamping force are fitted using single peak Gaussian function (black lines). Grey (RS), Red (pG) and blue (GG) colored histogram are indication that as the force increases the mean unfolding contour length shifts too.

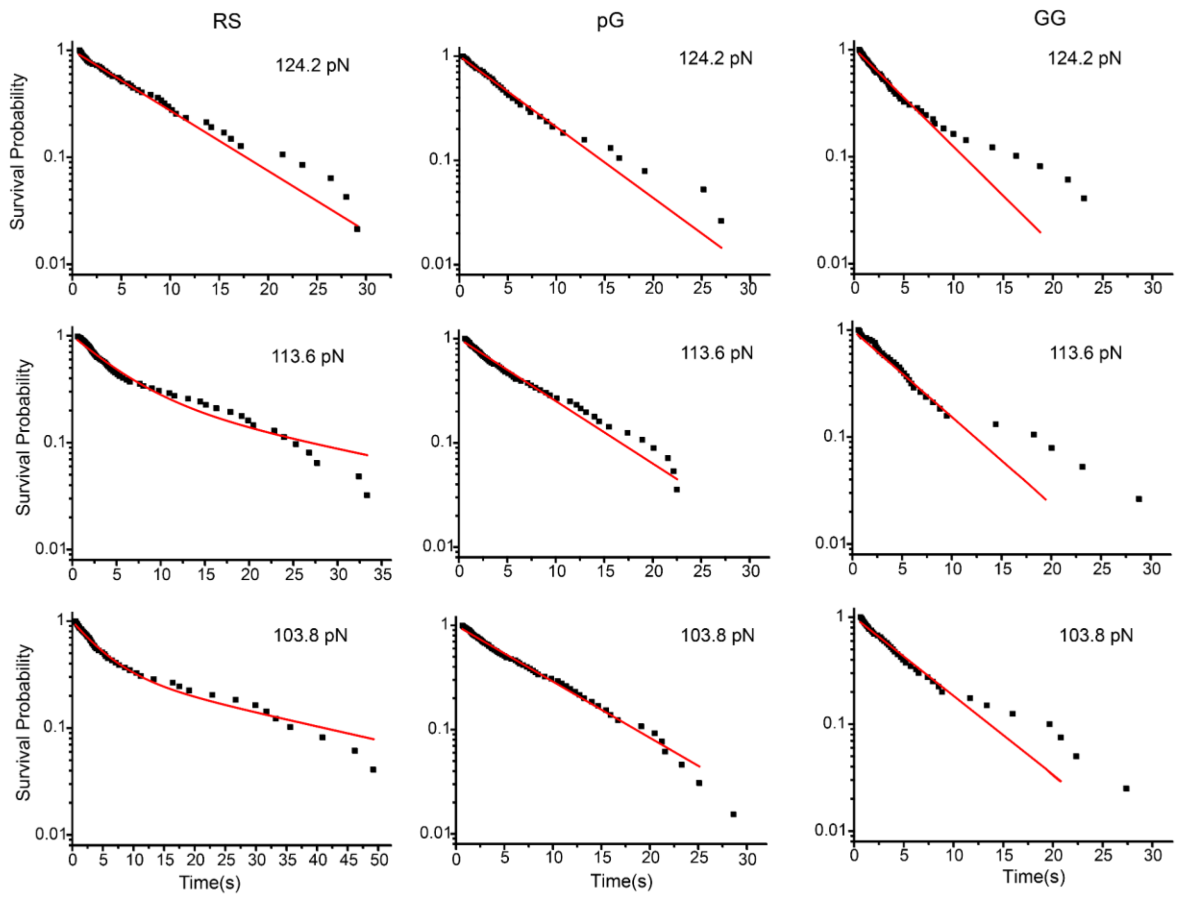


**SF 4: Survival probability plots for RS, pG and GG linker variants at different clamping forces from folded state:** The exponential decay fit ($y = \sum_{i=1}^{N} A_i * \exp(-x/\tau_i)$) of the survival plot at each clamping force is shown separately. The unfolding of I27 doublet in presence of IDL RS, pG, GG at force 103.8pN, 113.6 pN, 124.2pN.

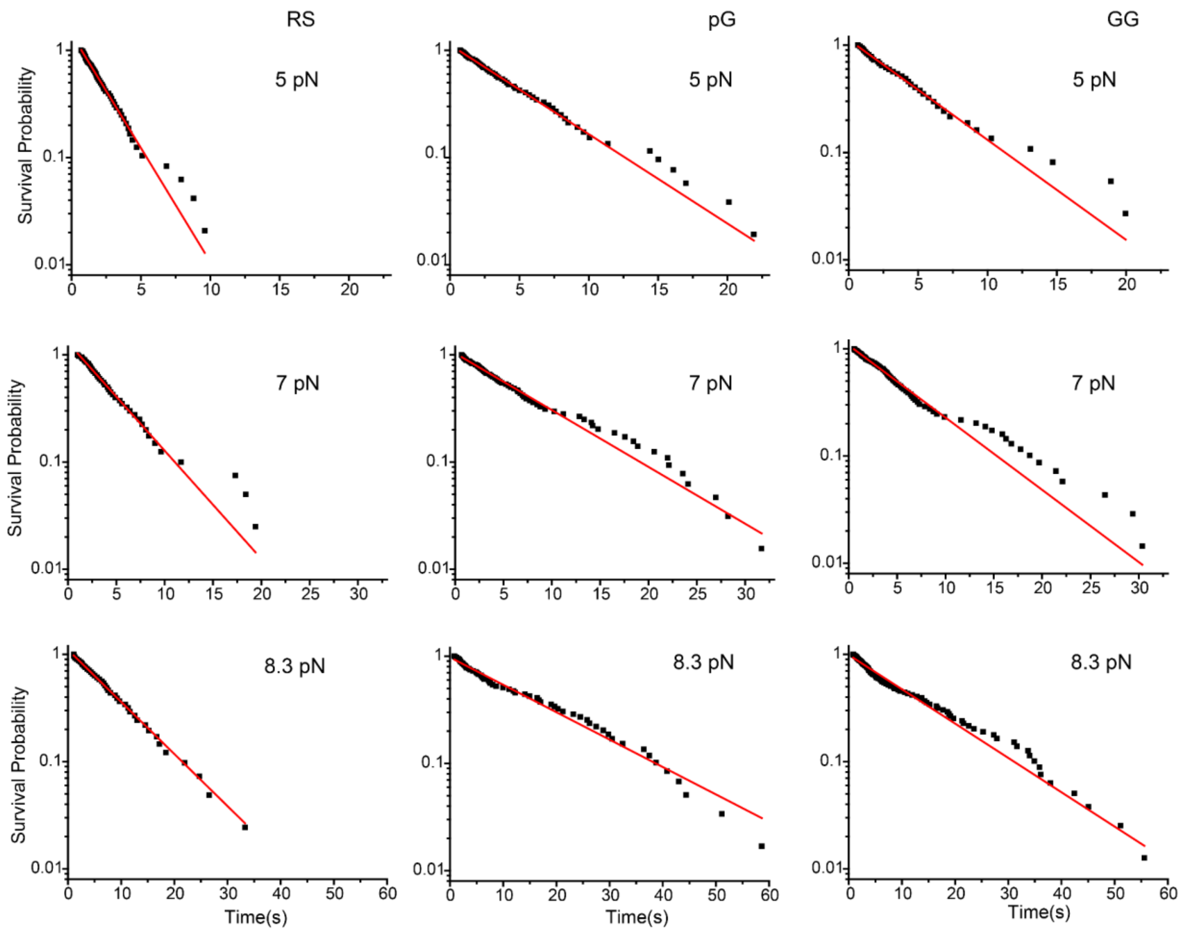


**SF 5: Survival probability plots for RS, pG and GG linker variants at different clamping forces from unfolded state:** Mono-exponential decay fit $(y = \sum_{i=1}^{N} A_i * \exp(-x/\tau_i))$ of the survival plot is shown separately at each clamping force of 5.0, 7.0 and 8.3 pN respectively. The refolding data of I27 doublets RS, pG, GG variants.

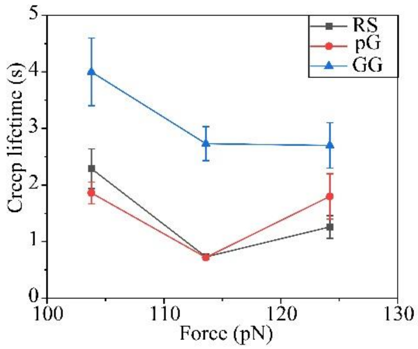


**SF 6: Rate of creep like transition for all three variants:** The lifetime of unfolding creep events at 103.8, 113.6 and 124.2 pN are measured from single exponential fitting of the dwell time data coming from creep events and shown using this dot plot for RS (in black), pG (in red) and GG (in blue). We observe that in GG the creep is slower for all forces and as we increase unfolding force, the creep lifetime decreases for RS compared to pG.

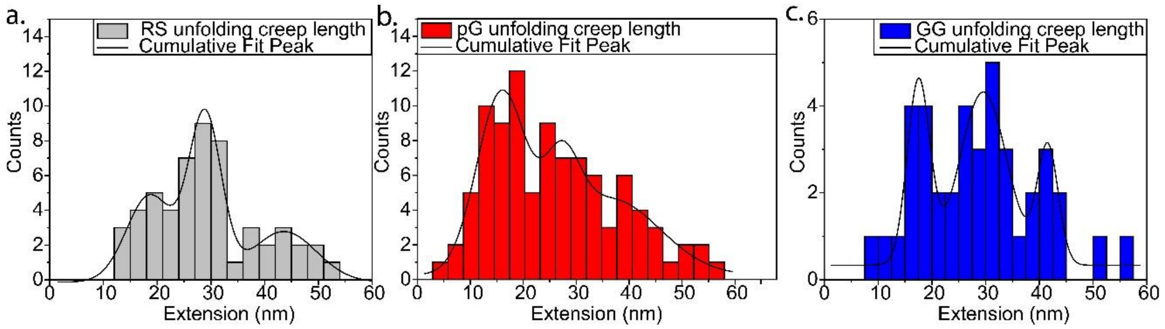


**SF 7**: **Creep length distribution of all three variants:** Distributions of unfolding creep lengths at forces, 103.8, 113.6, and 124.4 pN are plotted for RS (black), pG (red), and GG (blue) linker variants. The bin width was set to 3.0 nm for RS, 2.9 nm for pG, and 2.5 nm for GG to effectively capture creep patterns. The observed distribution reveals a strong correlation between creep length and expected contour length of individual domains, with a predominant peak in the range of ~25-30 nm, corresponding to one full domain unfolding, and a second peak below 20 nm corresponding to partial domain unfolding. Additionally, rare events exhibited extended creep length of ~40 nm, corresponding to more than one domain unfolding, indicating instances where two domains unfold in a sequential manner.

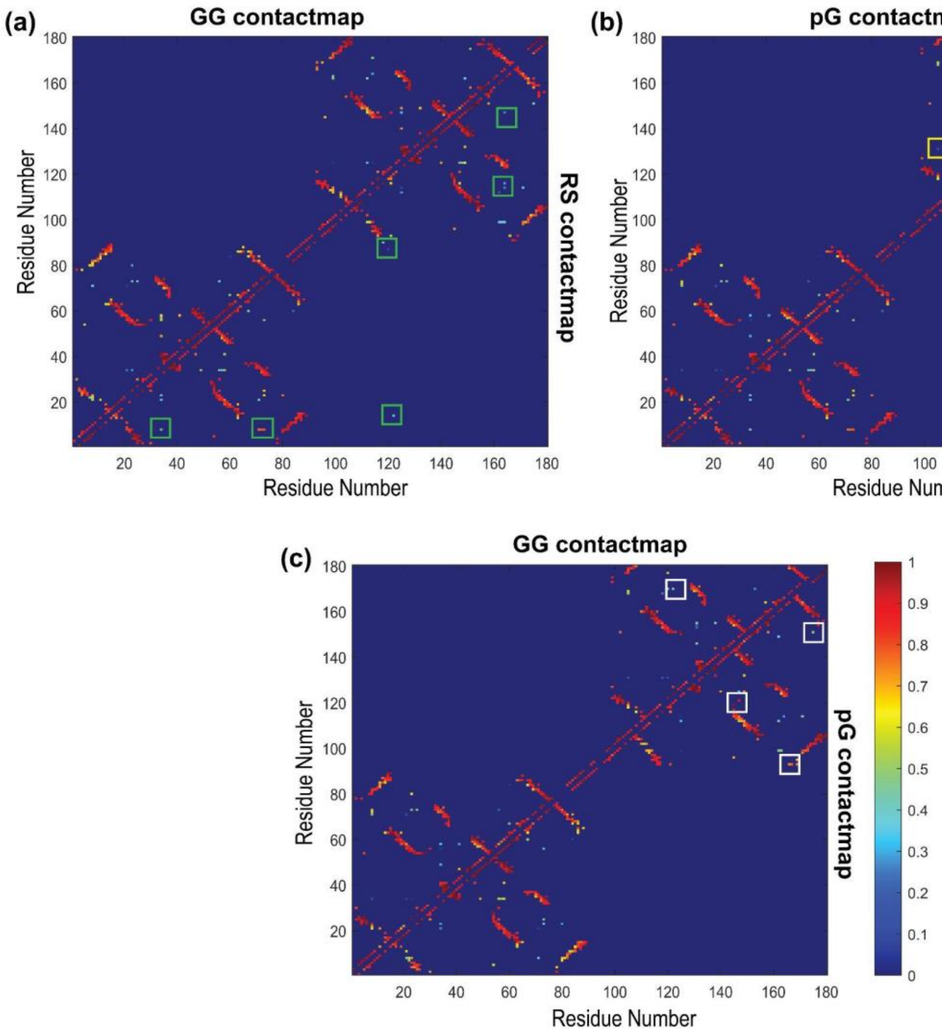


**SF 8: Graphical representation of the contacts or interactions between amino acid residues in RS, pG, and GG linker variants shown by contact map**: Contact map represents the close proximity of each amino acid in three-dimensional structure **(a)** The upper part of the contact map shows contacts for the GG linker variant, while the lower part shows those for the RS linker variant. Green squares highlight the differences in contacts between RS and GG linker variants. **(b)** The upper part of the contact map shows contacts for the pG linker variant, while the lower part shows those for the RS linker variant. Yellow squares highlight the contact differences between two. **(c)** Similar to **(a)** and **(b),** white squares in **(c)** show contact differences between GG and pG linker variants. Contacts within a certain distance range are colored differently to show strong versus weak interactions. Strong contacts (red cells in the map) indicate stable interactions, while weaker contacts (blue cells) indicate more transient interactions or flexibility in the structure.

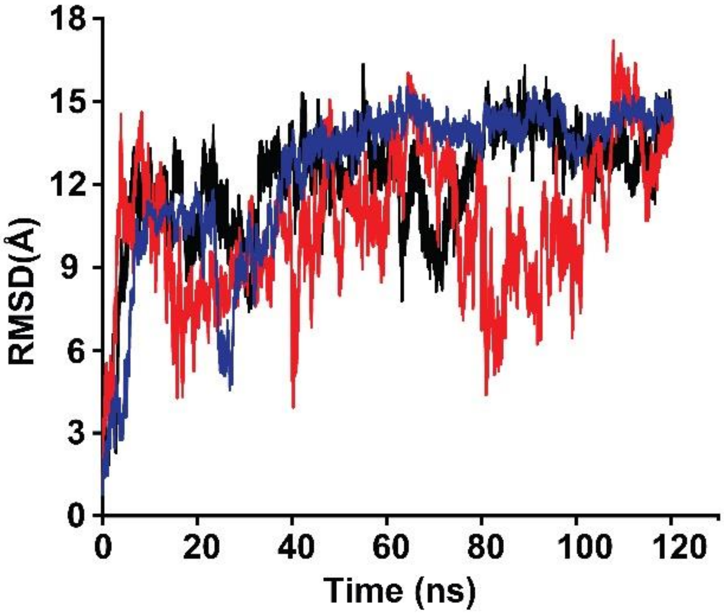


**SF 9: RMSD plot for RS (black), GG (blue), and pG (red) variants: A comparison of structural dynamics.** Average RMSD plots for RS, GG, and pG variants over 120 ns simulation time. Two independent simulations were performed for each protein, and the average RMSD values are shown. The plot indicates that the RMSD of pG variants fluctuating over the simulation time, a more flexible structure while the RS, GG variants display a stable RMSD after a certain time indicate that their structure remains relatively unchanged in simulation.

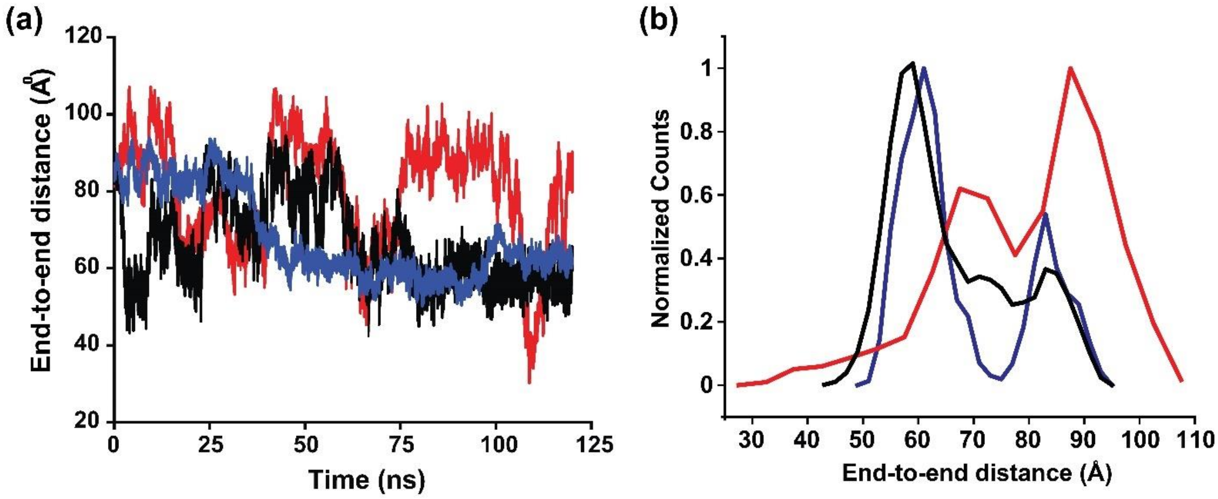


**SF10: Conformational dynamics of RS (black), GG (blue), and pG (red) variants: Insight from end-to-end analysis. (a)** End-to-end distance profile for RS, GG, and pG variants over 120 ns simulation. The profile shows that pG has higher and flexible end-to-end distance compared to RS and GG variants, indicate pG adopt a more extended and dynamic conformation, whereas RS and GG variants maintain a compact and stable conformation. **(b)** Counts vs end-to-end distance plot**.**

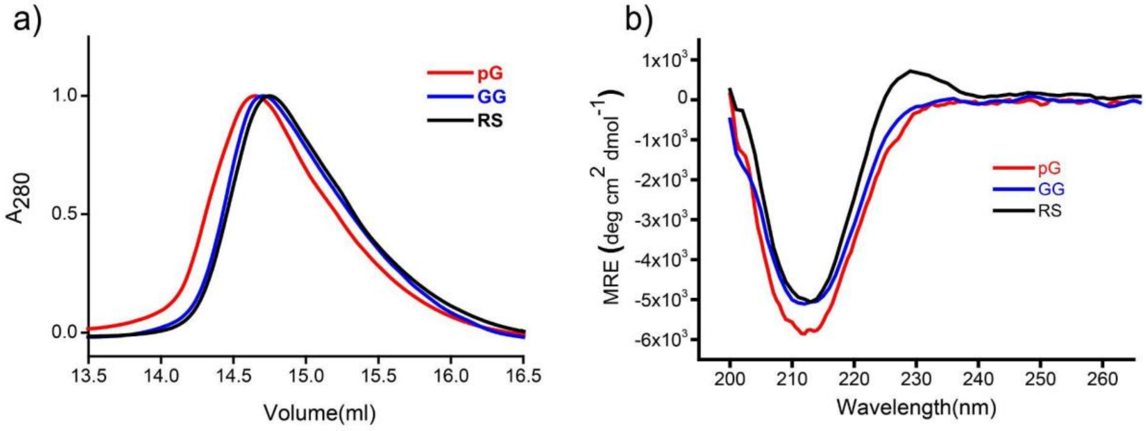


**SF 11: Comparative size-exclusion chromatogram (SEC) of the I27 doublet variants monitored using absorbance at 280 nm.** In SEC molecules elute according to their hydrodynamic volumes, suggesting equal molecular size of (I27-RS-I27) and (I27-GG-I27) whereas (I27-pG-I27) is slightly larger. Similar Rg for $(I27)_2$RS and $(I27)_2$GG indicates similar conformation.

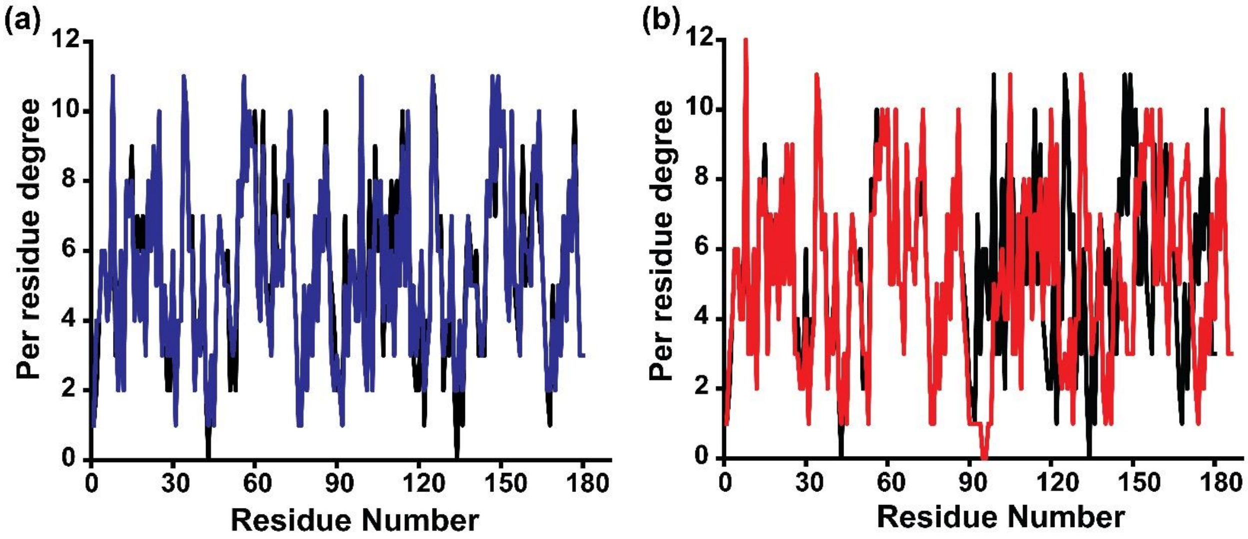


**SF 12: Comparison of residue interactions (degree analysis) in RS, Pg, GG linker variants with respect to RS variant. (a)** Degree comparison between RS and GG linker variants highlighting similarities and differences in their residue interaction patterns. **(b)** Degree comparison between RS and pG linker vaiants highlighting interaction profile between these two proteins. In protein interaction network, degree of each residues represents the number of interactions formed with the neighboring residues. This data suggests that linkers do not change the interaction pattern within the domains. Although some visual differences are there in figure b, that is due to the shift of amino acid position due to long linker (pG).

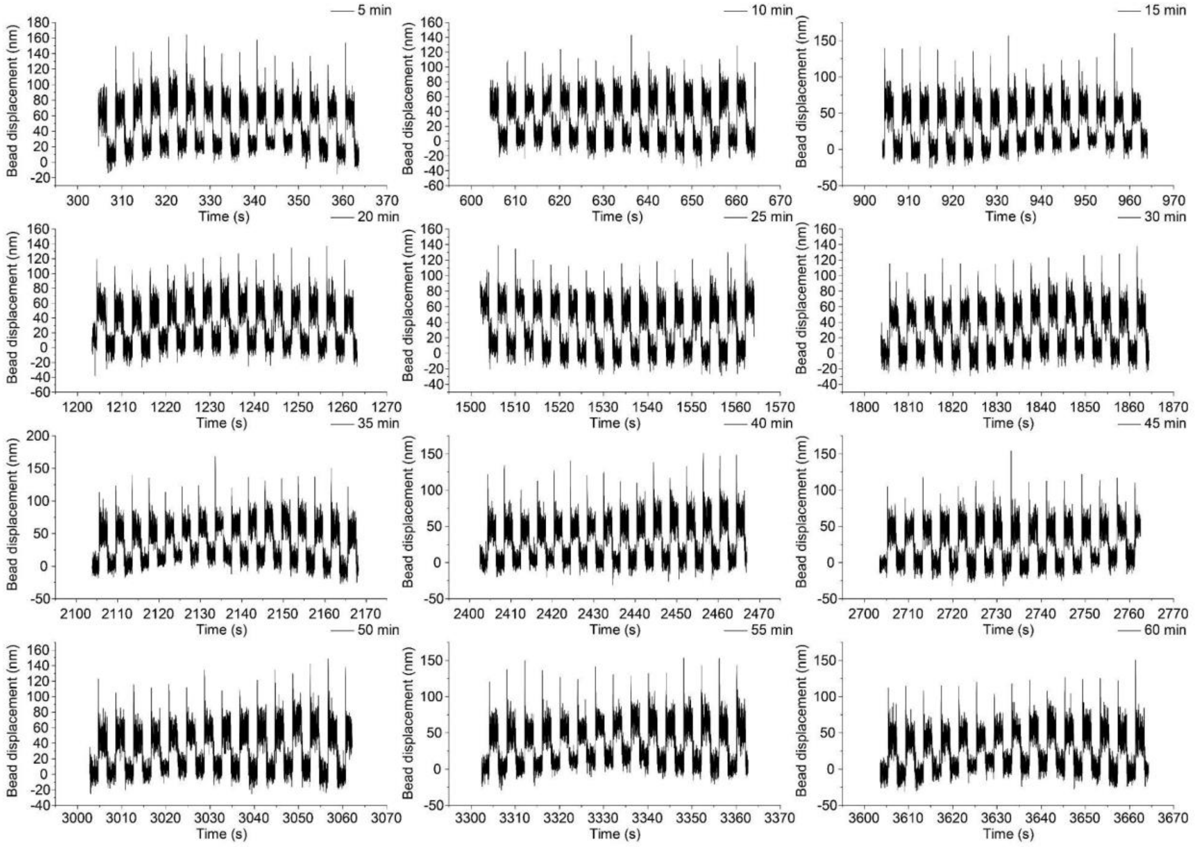


**SF 13**: **Plot of 60-second traces recorded at 200 Hz frame per seconds after every 300 seconds while the beads are subjected to oscillatory force-pulse of 0.25 Hz.** These are bead position (nm) versus time (s) trace data.

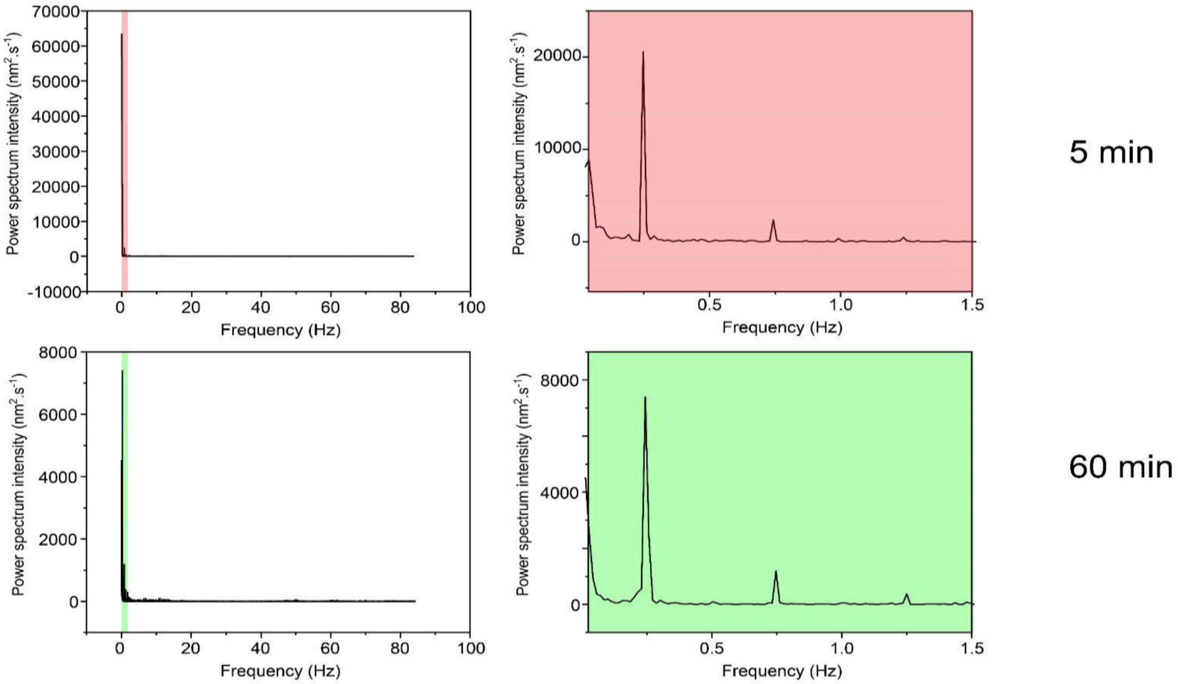


**SF 14: Power spectrum analysis example:** Power spectrum plot of 60 second recorded data of 305-365 seconds (top left plot) and 3605-3665 seconds (bottom left plot) showing intensity (nm2.s-1) versus frequency (Hz). The region from 0.1- 1.5 Hz has been highlighted in both plots and zoomed in versions have been plotted on the side panel. Pink box (power spectrum of extension vs time trace data recorded in between 305-365 seconds) shows the higher intensity at 0.25 Hz than intensity at 0.25 Hz of green box (power spectrum of extension vs time-trace data recorded in between 3605-3665 seconds) suggesting with time the power output of proteins decreases when the tether is experiencing continuous force pulse.

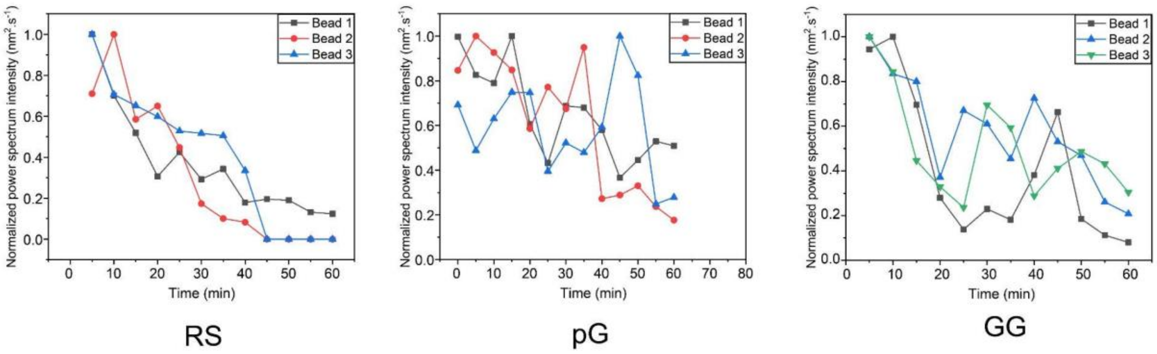


**SF 15: Decrease in power with time for all three variants.** Power spectrum analysis of 60 seconds long extension versus time trace data, is done for each of the one-minute time trace data which is collected after every five minutes and a normalized average intensity is plotted with respect to time for all three variants. The number of beads for this experiment was n=3 for pG, n=6 for GG and n=3 for RS. Three beads are shown with three different colors: black, red and blue. The average with error bars as standard deviation of these three values were plotted in Figure 4d.

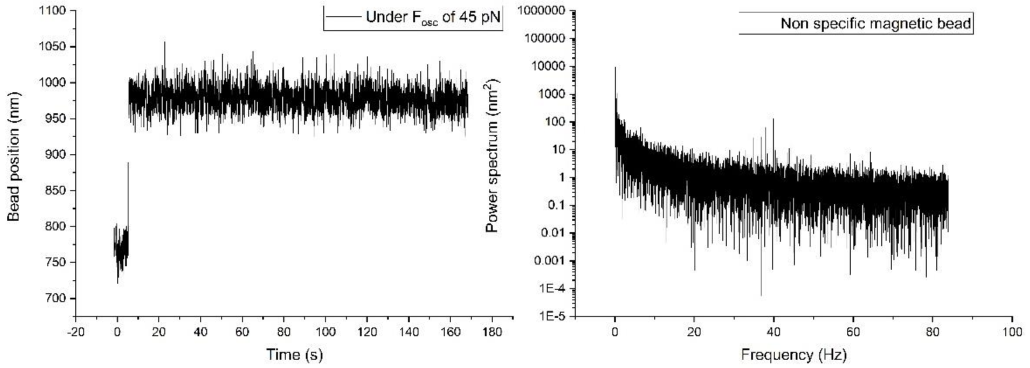


**SF16: Example of data and power spectrum analysis from Non-specific magnetic bead.** These beads do not oscillate. There is no peak in the power spectrum. No peak is being observed around 0.25 Hz, thus there is no signal and no decay with time or damping.
